# Neural investigation of idea selection during creative thinking reveals value-based mechanisms

**DOI:** 10.64898/2026.03.06.710072

**Authors:** Sarah Moreno-Rodriguez, Benoît Béranger, Emmanuelle Volle, Alizée Lopez-Persem

**Affiliations:** FrontLab, Sorbonne University, Institut du Cerveau - Paris Brain Institute - ICM, Inserm, CNRS, AP-HP, Hôpital de la Pitié Salpêtrière, Paris, France; CENIR, Institut du Cerveau - Paris Brain Institute - ICM, Inserm, CNRS, AP-HP, Hôpital de la Pitié Salpêtrière, Paris, France

## Abstract

Creative thinking enables humans to innovate and solve problems, via the generation of candidate ideas, their evaluation and the selection of the most promising one. While recent studies have demonstrated that idea evaluation relies on the assignment of subjective values (the assessment of how much one likes an idea, or valuation), the selection mechanism remains unknown. Using behavioral experiments and fMRI, we tested whether the selection step of creative thinking is driven by the same value comparison circuits that guide economic decisions. Participants performed creative idea production, rating, and choice tasks. We found behavioral evidence of value-based comparison during both isolated choice and overall creative idea production, in a manner that predicts creative abilities. Critically, we demonstrate that creative idea selection involves the brain valuation system and the dorsal anterior cingulate cortex: during creative thinking, these neural circuits compare candidate idea values, mirroring economic choice. Our findings reveal that idea selection is mediated by the brain’s generic valuation system, rather than specialized creative machinery, unifying theories of decision-making and creativity.

## 1. Introduction

Creative thinking is a cognitive function that enables humans to innovate and solve problems, via the production of creative ideas. In neuroscience, a creative idea is consensually defined as both original (or infrequent, unusual, novel) and adequate (or useful, appropriate, effective) (Runco & Jaeger, 2012). To produce such an idea, individuals must go through multiple thinking stages, starting with the generation of candidate ideas, followed by their evaluation, leading to the selection of the best idea, out of all the candidates.

At the cognitive level, idea generation is thought to rely on the novel combination of remote concepts stored in memory. This stage has long been considered to rely on spontaneous processes (Sowden et al., 2015), such as associative mechanisms (Beaty & Kenett, 2023; Mednick, 1962), but growing evidence suggests that controlled processes may also be involved and play a guiding role in idea generation (Altmayer et al., 2025; Volle, 2018). In contrast, idea evaluation is thought to rely on the assessment of the idea’s originality and adequacy, though few studies have directly demonstrated that these dimensions are actually considered (Moreno-Rodriguez et al., 2025). Idea evaluation has traditionally been associated with controlled processes (Sowden et al., 2015; Wang et al., 2024), such as memory-based assessment (Benedek et al., 2023) and the suppression of distractions (Chrysikou, 2019). More recently, idea evaluation has been associated with subjective valuation, i.e. how much one *likes* an idea, a process known to underlie evaluation, but also to guide selection, in value-based decision-making research (Kleinmintz et al., 2019; Lin & Vartanian, 2018; Lopez-Persem et al., 2024; Moreno-Rodriguez et al., 2025).

At the neural level, creative idea generation has been consistently associated with the Default Mode Network (DMN): a network comprising the medial prefrontal cortex, posterior cingulate cortex, precuneus, inferior parietal lobule, and temporal regions. It is known to support associative thinking, mind-wandering, and overall internally-oriented cognition (Zabelina & Andrews-Hanna, 2016). In parallel, idea evaluation has been consistently associated with the Executive Control Network (ECN): a network comprising the dorsolateral prefrontal cortex, anterior cingulate cortex, inferior parietal lobule, and temporal areas. This network is typically associated with cognitive control processes (Niendam et al., 2012). Interestingly, a growing number of findings challenge the strict matching of the DMN and ECN onto generative and evaluative processes, instead suggesting that both networks are involved in both stages (Beaty et al., 2014; Ellamil et al., 2012; Lloyd-Cox, Chen, et al., 2022; Moreno-Rodriguez et al., 2025; Shi et al., 2018). In parallel, the salience network (SN) has been associated with regulating the balance between ECN and DMN activity (Beaty et al., 2015, 2016).

In summary, the cognitive and neural bases of the generation and evaluation steps of creative thinking have been largely explored. In contrast, the final selection step lacks a precise cognitive and neural account. So far, it has been mostly studied through the lens of selection accuracy, i.e., individuals’ ability to recognize the most creative idea among several options, as compared to the opinion of independent judges, usually creativity experts. A consistent finding in this line of research is that people are often suboptimal in their selection (Rietzschel et al., 2006, 2010; Zhu et al., 2017). Various selection strategies have been proposed to increase selection accuracy, such as changing individuals’ state of mind (de Buisonjé et al., 2017) or giving them specific selection instructions (Birney et al., 2016; Rietzschel et al., 2010; Zhu et al., 2021). Importantly, Zhu et al., (2021) found that following one’s intuition led to a more optimal selection than following specific instructions. This intuitive selection may relate to the personal attachment that individuals develop towards the ideas that they find “desirable”, as reported by several studies (de Buisonjé et al., 2017; Eling et al., 2015; Lazar et al., 2022; Zhu et al., 2021). Interestingly, this desirability could echo the role subjective valuation in creativity mentioned earlier (Kleinmintz et al., 2019; Lin & Vartanian, 2018; Lopez-Persem et al., 2024; Moreno-Rodriguez et al., 2025). Yet, not all studies find intuitive selection to be more accurate. Rietzschel et al. (2010) report that this process favours feasible and desirable ideas over original ones, and Lazar et al. (2022) argue that it can bias selection toward ideas aligned with an individual’s self-concept, which can easily diverge from experts opinions and thus lead to lower selection accuracy.

Overall, selection is the critical bottleneck that allows committing to a creative idea, yet it remains the least understood phase of creative cognition. While idea generation and evaluation have been mapped to distinct neural circuits, the mechanism by which the brain commits to a single idea has been mechanistically obscure, empirically confounded with evaluation and misattributed to domain-general executive control networks. This gap represents a fundamental barrier to understanding how humans solve problems. In the current study, aligning with Zhu et al. (2021)’s view on idea “desirability”, we argue that individuals select the idea they personally like the most. In effect, we propose that the selection step of creative thinking is guided by the comparison of the subjective values assigned to candidate ideas, similarly as during value-based decision-making.

Extensive research in value-based decision-making has demonstrated that when faced with a choice, options are assigned a subjective value, referred to as the valuation step. Then, options are compared through the computation of a decision value, i.e., the difference between two options’ values. This is referred to as the decision step. Finally, based on the result of this comparison, the option with the highest value is selected (Glimcher, 2014; Kahneman & Tversky, 1979; Samuelson, 1937).

In value-based decision-making research, subjective values can be measured using rating tasks or choice tasks (Lopez-Persem et al., 2017). During rating tasks, participants explicitly declare the value they assign to an option but they do not need to forgo one option in favor of another. In contrast, during choice tasks, participants face two (or more) options, and must select the one they prefer. By forcing participants to discriminate between options, choice tasks reveal preferences in an implicit manner. Choice tasks are also more ecologically valid, as everyday situations involve selecting between alternatives more often than explicitly rating them. Therefore, although both task types enable the capture of underlying hidden subjective values, they can yield subtly different outputs. For example, choices do not indicate the strength of preference, only relative preferences, but choice response times can be used as an informative continuous measure: since they decrease with increasing preference strength (De Martino et al., 2013; Konovalov & Krajbich, 2019), they can be used to rank preferences and predict whether individuals will stick with their choice or change their mind later. Additionally, choice tasks are better suited for uncovering cognitive biases (Lopez-Persem et al., 2016).

Value-based decision-making research has also identified brain regions supporting subjective valuation and comparison processes. First, subjective valuation has been associated with a number of regions referred to as the Brain Valuation System (BVS): a network that overlaps the reward system identified in animal literature, comprising notably the ventromedial prefrontal cortex (vmPFC) and the ventral striatum (Bartra et al., 2013). Importantly, previous work from our lab showed that the BVS also supports valuation during creative thinking (Moreno-Rodriguez et al., 2025). Then, subjective value comparison has been associated with both the BVS, typically encoding the decision value framed as the value of the chosen option minus the value of the unchosen option (Boorman et al., 2009; Kable & Glimcher, 2009; Rangel & Hare, 2010), and the dorsal anterior cingulate cortex (dACC), which negatively encodes decision values (Chong et al., 2017; Lopez-Persem et al., 2016; Wunderlich et al., 2009), probably reflecting accumulation processes and guiding selection (Basten et al., 2010; Hunt et al., 2014; Wunderlich et al., 2009). However, the exact computation occurring in this brain region is debated (Clairis & Lopez-Persem, 2023): other studies associate the dACC with the negative encoding of subjective values (Bartra et al., 2013), or with the tracking of conflict and difficulty signals (Gloy et al., 2020; Volz et al., 2005), expected value of control (i.e. the level of cognitive control required to complete a demanding task) (Shenhav et al., 2013; Shenhav & Botvinick, 2015), or the value of exploring alternatives in the exploration/exploitation framework (Boorman et al., 2013; Hayden et al., 2011; Kolling et al., 2012; Quilodran et al., 2008; Tsetsos et al., 2014). In any case, the BVS and the dACC emerge as key regions for studying the value-based comparison processes that guide option selection.

In summary, while selection processes have been thoroughly investigated in value-based decision-making research, idea selection processes have been overlooked in creativity research. In the current study, we conceptualize idea selection as an integral step of creative thinking and use a value-based framework and functional Magnetic Resonance Imaging to address this gap in the literature. In our framework (Lopez-Persem et al., 2024), we posit that creative thinking, after the generation of candidate ideas, relies on the same value-based decision-making steps as classical value-based decision making: individuals first assign them subjective values (valuation). Then, they compare the candidate ideas’ subjective values (comparison) to select the one they prefer (selection). In previous studies, we demonstrated that idea evaluation does indeed involve subjective valuation: during a creativity task, individuals assigned subjective values to ideas (Lopez-Persem et al., 2024). Using neuroimaging, we identified the neural correlates of this subjective valuation of ideas, finding that, similarly to the subjective valuation of food or other types of items, ideas’ values were encoded in the BVS (Moreno-Rodriguez et al., 2025). Then, Lopez-Persem et al. (2024) went further and demonstrated that idea selection involves a comparison of these subjective values: during a creativity task, individuals’ responses could be predicted by a model that compared the ideas’ subjective values and selected the idea with the highest value. Using neuroimaging, we now aim to reveal the neural correlates of the value-based comparison of ideas. Based on the decision-making literature, we hypothesize that these neural correlates will involve the BVS and the dACC region.

To test this hypothesis, 108 participants performed a creative idea production task. Then, they evaluated the likeability, originality and adequacy of their ideas during rating tasks. Finally, they selected their preferred ideas during a choice task. We first found that the decision values computed using the rating task differed from those computed using the choice task. Since the latter better explained individual choices, we used these decision values in the subsequent analyses investigating idea selection. Then, we found interindividual differences in the valuation of ideas during the choice task and showed that they related to interindividual differences in creative abilities, suggesting a link between value-based decision-making mechanisms and creative performance. Next, we regressed response times against decision values during the choice task and during the idea production task and found that in both tasks, response times decreased with decision values, suggesting that creative thinking involves the computation of a decision value before committing to a response. Finally, using fMRI in a subset of 38 participants, we identified a neural signature of value-based comparison of ideas during the choice task in the vmPFC and the dACC, and showed that this signature was also present during the creative idea production task, thus supporting the hypothesis that, during creative thinking, idea selection relies on the value-based comparison of ideas.

## 2. Results

The present study merges the data from two subgroups of participants who performed the same tasks in the same order. The first subgroup (subgroup A, n=69) performed all the tasks in front of a computer, while the second subgroup (subgroup B, n=39) performed some of them in an MRI scanner (Figure 1). They performed a creative idea production task (the Free Generation of Associates Task or FGAT, Figure 1B), a likeability rating task (Figure 1C), a choice task (Figure 1D), an originality and adequacy rating task (Figure 1E), and finally a battery of creativity tests.

**Figure 1:**
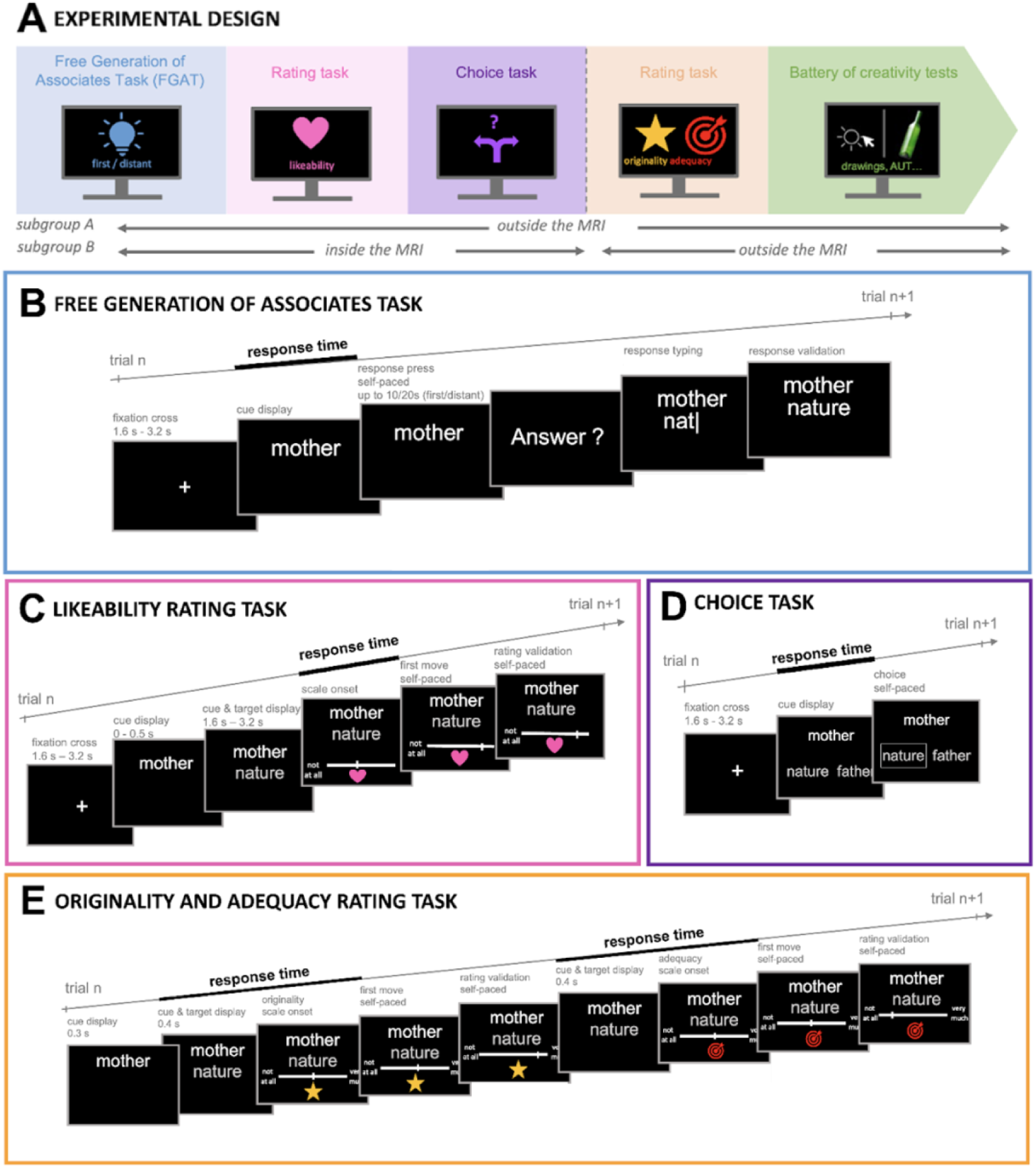
Experimental design. (A) Overview of the different tasks in chronological order (B) Free Generation of Associates Task (FGAT). In two conditions, participants had to provide: the first word that came to their mind when they saw the cue word (FGAT-first) or a more distant word, with the instruction to be creative (FGAT-distant). Participants outside the MRI (subgroup A) typed their answer themselves. Participants in the MRI (subgroup B) provided their responses orally to the experimenter, who typed them. They had the opportunity to repeat their response in case the experimenter had misheard. (C) Likeability rating task. Participants rated each association on a likeability scale from “not at all” to “very much”. (D) Choice task. Participants saw a cue word above two possible responses, and were instructed to choose the one they preferred. (E) Originality and adequacy rating task. Participants rated each association on a scale going from “not at all” to “very much”. In the MRI, all trials of all tasks began with a fixation cross, followed by the trial screen.

The FGAT is a remote thinking task reported to capture critical aspects of creativity (Battistello et al., 2025; Bendetowicz et al., 2018; Lopez-Persem et al., 2024; Moreno-Rodriguez et al., 2025; Prabhakaran et al., 2014). In the FGAT-first condition, considered a control condition, participants were presented with a cue word and had to respond with the first word that came to mind when reading it. Next, the FGAT-distant condition used the same cue words, but this time participants had to respond with a word that was further from the cue, with the instruction to think creatively (see Methods for more details).

In the rating tasks, participants saw FGAT cue-response associations, including their own FGAT-first and FGAT-distant responses, as well as FGAT-first and FGAT-distant responses from the database of a previous study of the lab that used the same FGAT (Altmayer et al., 2025). Participants rated how much they liked the cue-response associations (likeability ratings) and, later, how original and adequate they found them (originality and adequacy ratings), using pseudo-continuous scales going from ‘not at all’ to ‘very much’ (see Methods for more details).

In the binary choice task, each trial showed participants a cue word above two possible responses. Choice trials included all the different types of pairings between FGAT-first and FGAT-distant responses from the participants’ own responses and the previous study’s database (first/distant × participant/previous database). At each trial, participants had to choose the response they would have preferred to provide in the FGAT-distant condition (see Methods for more details).

### 2.1. Estimating subjective values used during value-based selection of creative ideas

In value-based decision-making, a choice is assumed to depend on a decision value (DV), defined as the difference between the subjective values assigned to available options. The greater this difference in favor of one option, the faster and more probable its selection. Thus, to investigate idea selection during creative thinking, we used a binary choice task to isolate decision-making mechanisms driving choices between creative ideas.

We first needed to determine which subjective values drive these choices. Previous work that used likeability ratings has shown that subjective value attributed to an idea relies on a combination of the idea’s adequacy and originality (Battistello et al., 2025; Lopez-Persem et al., 2024; Moreno-Rodriguez et al., 2025) (see Equation B in Methods). Nevertheless, previous research suggests that subjective values estimated from likeability rating tasks can slightly differ from those inferred from choices (Lopez-Persem et al., 2016; Sullivan & Huettel, 2021). We must therefore consider two distinct mechanisms for DV computation, which could result from either a “value-before-choice” mechanism, during which individuals compute subjective values similarly as in a likeability rating task *and then* compare them to make choices (thus, DVs can be estimated via likeability ratings); or a “value-during-choice” mechanism, during which individuals assign values to options *while* they are comparing them to make choices (thus DVs can be estimated directly from choices).

To clarify which of these two valuation mechanisms better explain the choices of creative ideas, we used a computational modelling approach (see details in Methods and Supplementary Material) to compare two types of models. The “value-before-choice” models first fitted a value function to likeability ratings (with adequacy and originality ratings as inputs), and these values were then used to predict choices in a softmax function (see model equations in Methods). On the other hand, the “value-during-choice” models directly fitted a softmax function including a value function (with adequacy and originality ratings as inputs) to choices, such that subjective values were estimated directly from choices (see model equations in Methods).

We found that the model that best explained choices was a “value-during-choice” model (see Equation C in Methods) (Figure 2A & B). This model included the parameters *⍺*_choice_ and *δ*_choice_, hereafter referred to as valuation parameters, that capture how individuals weigh originality and adequacy during the choice task. The estimation of the *⍺*_choice_ parameter revealed that, on average, participants put less weight on originality than on adequacy when comparing creative ideas. The estimation of the *δ*_choice_ parameter showed they had a preference for ideas showing an equilibrium of originality and adequacy over ideas that are more extreme on one or the other dimension (see Table S2).

**Figure 2:**
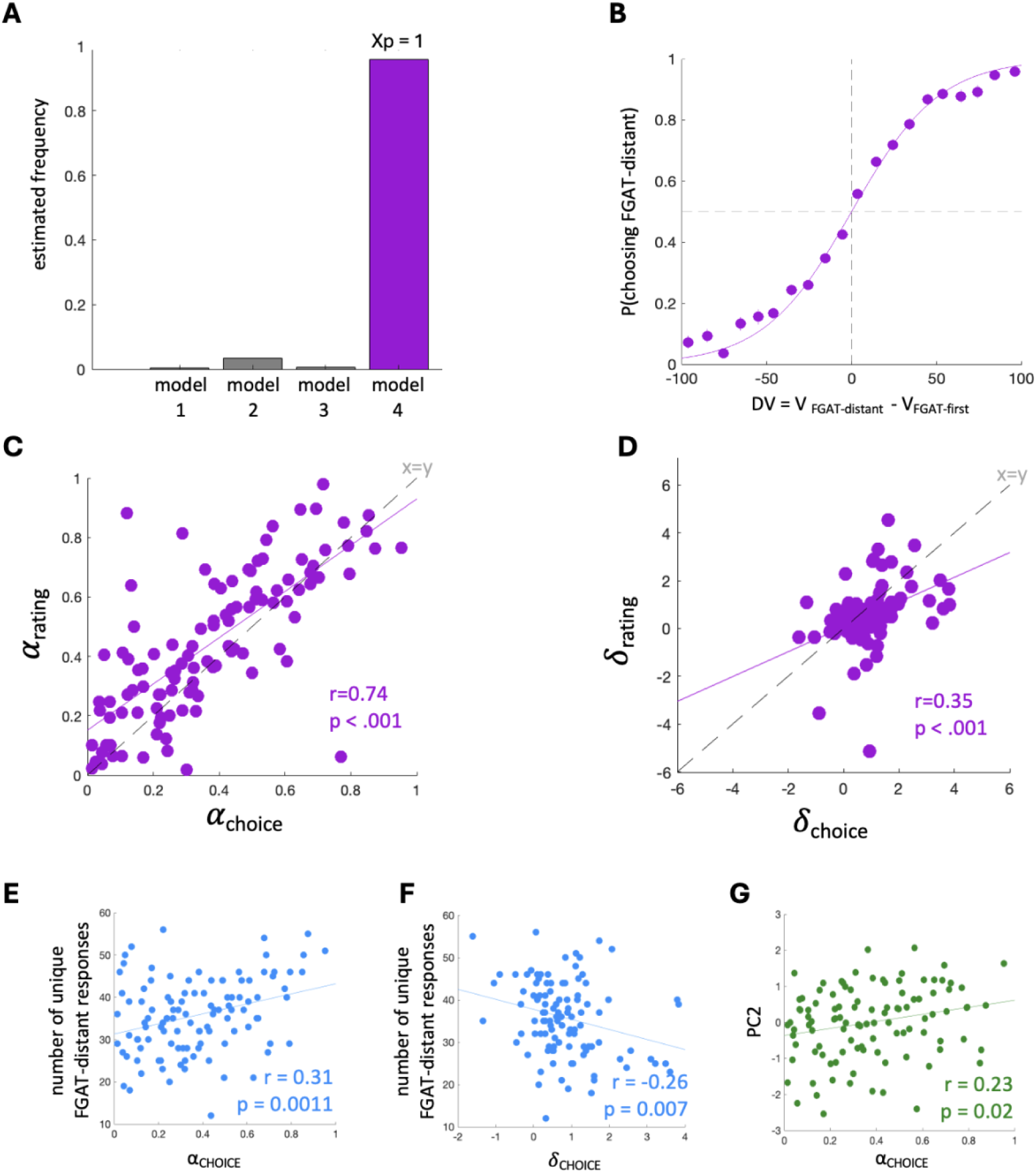
Model comparison and valuation parameters estimation linking choice behavior and creative performance. **(A)** Estimated frequencies of the four models included in the model comparison and exceedance probability (Xp) of the winning model (model 4). Model equations can be found in the Methods. **(B)** Model fit of model 4 on the choice task data: the probability of choosing the FGAT-distant option can be explained as a softmax function of the decision value (DV, the difference of values between the FGAT-distant and the FGAT-first response). Data points indicate binned data averaged across participants. Error bars are intersubject SEM. **(C)** Correlation between α_choice_ and α_rating_. Each data point corresponds to one participant. The continuous purple line indicates a significant correlation (p<0.001). The grey dotted line is the identity function. The majority of data points above the identity (72 out of 108) corresponds to participants who lowered their α (i.e., their weighting of originality) during choices compared to ratings (and vice-versa). **(D)** Correlation between *δ*_choice_ and *δ*_rating_. Each data point corresponds to one participant. The continuous purple line indicates a significant correlation (p<0.001). The grey dotted line is the identity function. The majority of data points below the identity (63 out of 108) corresponds to participants who increased their *δ* (i.e., decreased their preference for a balance of originality and adequacy) during choices compared to ratings (and vice-versa). **(E)** Correlation between the *⍺*_choice_ parameter and the number of unique associations in the FGAT-distant. **(F)** Correlation between the *δ*_choice_ parameter and the performance in the FGAT-distant. **(G)** Correlation between the *⍺*_choice_ parameter and the second PCA component (PC2). The solid lines indicate significant correlations (p<0.05).

Then, we compared the valuation parameters from the winning “value-during-choice” model (*⍺*_choice_ and *δ*_choice_) to those of the “value-before-choice” model (*⍺*_rating_ and *δ*_rating_). We found that *⍺*_choice_ was lower than *⍺*_rating_ and that *δ*_choice_ was lower than *δ*_rating_ (Figure 2C & D, Supplementary Results and Table S2), indicating that, during the choice task, individuals’ weighting of originality and preference for equilibrium had both dropped, compared to during the likeability rating task. Yet, they correlated between tasks, suggesting that, despite slight differences, preferences stayed coherent within individuals (Figure 2C & D, Supplementary Results and Table S2).

Additionally, we sought to confirm that the winning “value-during-choice” model (which used α_choice_ and *δ*_choice_ as valuation parameters) was relevant to creative performance. Using the number of unique answers in FGAT-distant and the battery of external creativity tests (see Methods for details), we found that participants favoring originality during creative idea comparison (higher α_choice_) yielded higher creative performance (Figure 2E-G, Supplementary Results and Table S2).

Finally, the winning “value-during-choice” model included two other parameters (see Equation C in Methods). A *β* parameter, (which captures the expected stochasticity, or incoherence, in individuals’ choices) and a *γ* parameter, which revealed an additive bias for FGAT-distant ideas, against FGAT-first ideas (see Supplementary Results). In other words, FGAT-distant responses typically received a small value bonus when compared to FGAT-first responses. This aligns with the choice task instructions (i.e., to choose the response they would have preferred to provide in the FGAT-distant condition).

Overall, the model comparison and the parameter estimations informed us on how originality and adequacy are integrated into a subjective value and how they drive the selection of ideas. They revealed that the decision values computed from choices differed from the difference of subjective values computed from likeability ratings, and that the former predicted choices better than the latter. Consequently, we used the decision values computed from choices when looking for a behavioral and neural signature of value-based selection during creative idea production (section 2.3).

### 2.2. Behavioral evidence that creative idea production relies on the comparison of ideas’ values

We aimed to identify a behavioral signature of the comparison process using DV, in the choice task, but also in the idea production task. In the choice task, we simply defined decision values as DV = V_chosen_-V_unchosen_, where V is computed using originality and adequacy ratings, α_choice_ and *δ*_choice,_ similarly to model 4 in section 2.1. When regressing choice response times (RT_choice_) against DV, we observed a linear decrease of RT_choice_ with DV (β = −0.14±0.01, M±SEM; one-sample two-tailed t-test: t(107) = - 12.5, p < 0.001, Figure 3A). This result is consistent with value-based decision making, and suggests that idea selection involves a comparison of the subjective values of the two options, indicating that selection is a value-based process.

**Figure 3:**
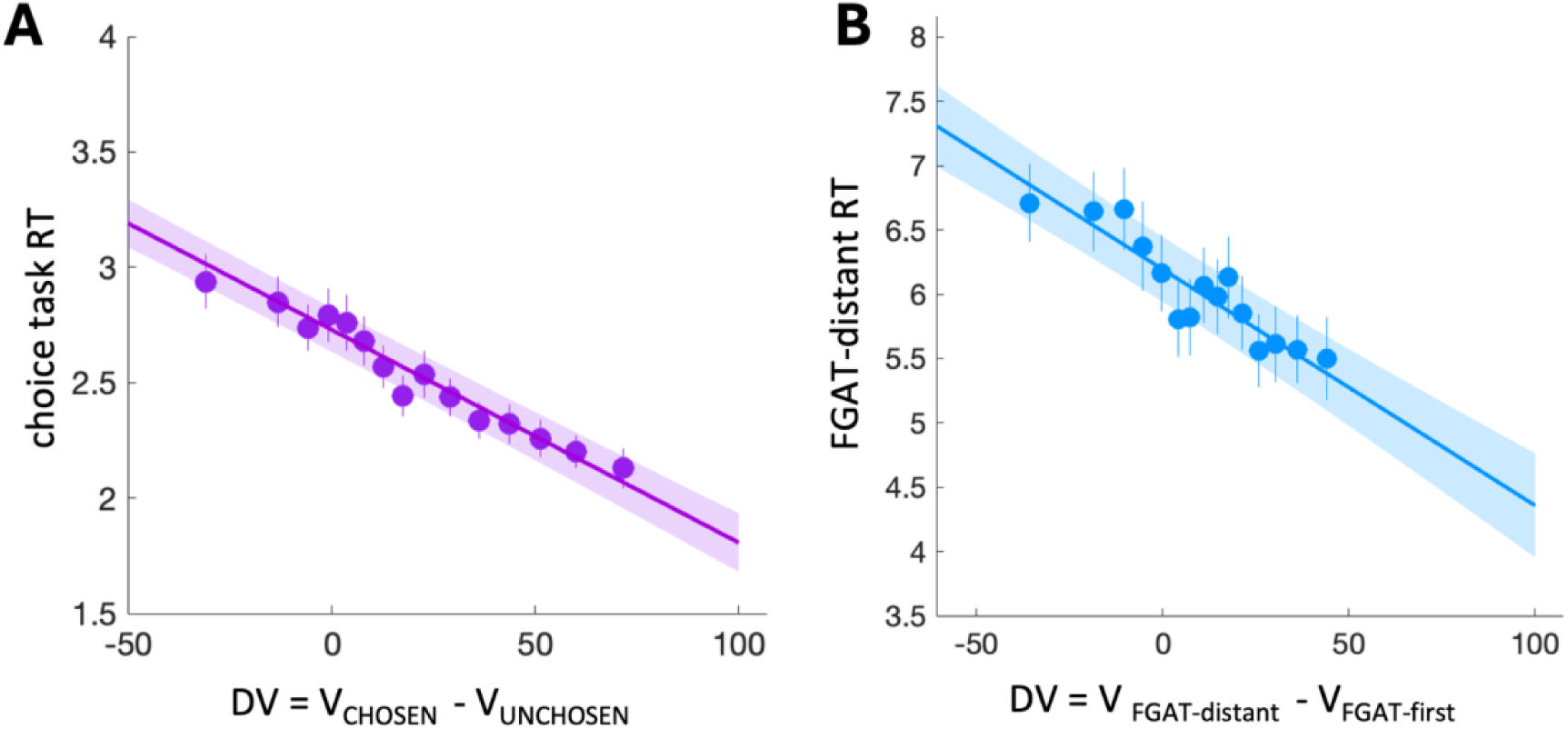
Behavioral evidence that creative idea production relies on the comparison of ideas’ values. (A) Model fit of model 4 on the choice task data: the probability of choosing the FGAT-first option can be explained as a softmax function of the decision value (DV, the difference of values between the FGAT-distant and the FGAT-first response). (B) Linear regression of the choice task response times (RTs) by the decision value (DV, the difference of values between the chosen and the unchosen response). (C) Linear regression of the FGAT-distant response times (RTs) by the decision value (DV, the difference of values between the FGAT-distant and the FGAT-first response). RTs and DVs in panels B and C are z-scored to account for intersubject variability. In all panels, data points indicate binned data averaged across participants. Error bars and shaded areas are intersubject SEM.

Then we looked for the same behavioral signature of the comparison process in the creative idea production task (FGAT-distant). While we propose that this task involves the value-based comparison of different ideas before one is chosen, our experimental design gives us access to only one alternative (unchosen) response per cue: the first response that comes to mind, the one that was given in the FGAT-first condition. Therefore, we considered that, during the FGAT-distant condition, the FGAT-first response to a cue is the unchosen option, while the FGAT-distant response to the same cue is the chosen option. Thus, we computed decision values as DV = V_FGAT-distant_-V_FGAT-first_. Again, V is computed using originality and adequacy ratings and α_choice_ and *δ*_choice_, similarly to the model 4 of section 2.1. Note that using FGAT-first responses as alternative ideas is not ideal since the first response might be quickly discarded during the search for a creative response, but it still provides a useful opportunity to investigate whether and how a comparison process is involved in creative thinking. Thus, we regressed RT_FGAT-distant_ against DV and observed a pattern similar to that in the choice task (β = −0.11±0.02, M±SEM; one-sample two-tailed t-test: t(107) = −6.11, p < 0.001, Figure 3B), indicating that responses are slower when the alternative option’s value is close in value to the chosen option’s value (small DV). This result suggests that idea production involves a comparison of the subjective values of the actual response and an alternative response, indicating a value-based selection process.

As a control analysis, we aimed to confirm that FGAT-distant indeed involved a value-based comparison process by assessing whether RTs were better explained by DVs than by the value of the selected response (FGAT-distant) alone. In line with our hypothesis, model comparisons showed that RT_FGAT-distant_ were better explained by DV than by the value of the FGAT-distant response alone, suggesting that the value of the alternative (FGAT-first) also contributed to explain some variance in the RTs during the production of creative ideas (see Supplementary Results).

Overall the relationship between RTs and DVs during the choice task and the creative idea production task offer behavioral evidence that a value-based comparison process is at play during creative thinking.

### 2.4. Neural evidence that creative idea production relies on the comparison of ideas’ values

#### 2.4.1. The vmPFC and the dACC encode decision values during the selection of externally presented ideas

To determine whether and where DVs are computed in the brain during creative thinking, we first explored the neural representation of DVs during the choice task. Indeed, this served as a localizer in order to define the brain regions of interest (ROIs) encoding DVs during idea selection. For these analyses, similarly to the previous RT_choice_ analyses, DVs were computed as the difference of values between the chosen and the unchosen responses of the choice task.

To investigate the neural representation of DVs during idea selection, we analyzed the choice task using a whole-brain parametric modulation approach. We tested a Generalized Linear Model (GLM1) containing the DVs as a parametric modulator of a categorical boxcar regressor between the onset of the trial (i.e., the display of the cue word and the two possible responses) and the button-press to choose the response. We found a significant positive effect of DVs in the vmPFC (Figure 4A): (peak Montreal Neurological Institute (MNI) coordinates: [x = 6 y = 62 z = −20], one-sample two-tailed t-test: t(37) = 5.24, pFWE = 0.010, see Table 1 for all significant clusters), a region known to encode DVs in decision-making (Hunt et al., 2012).

**Figure 4:**
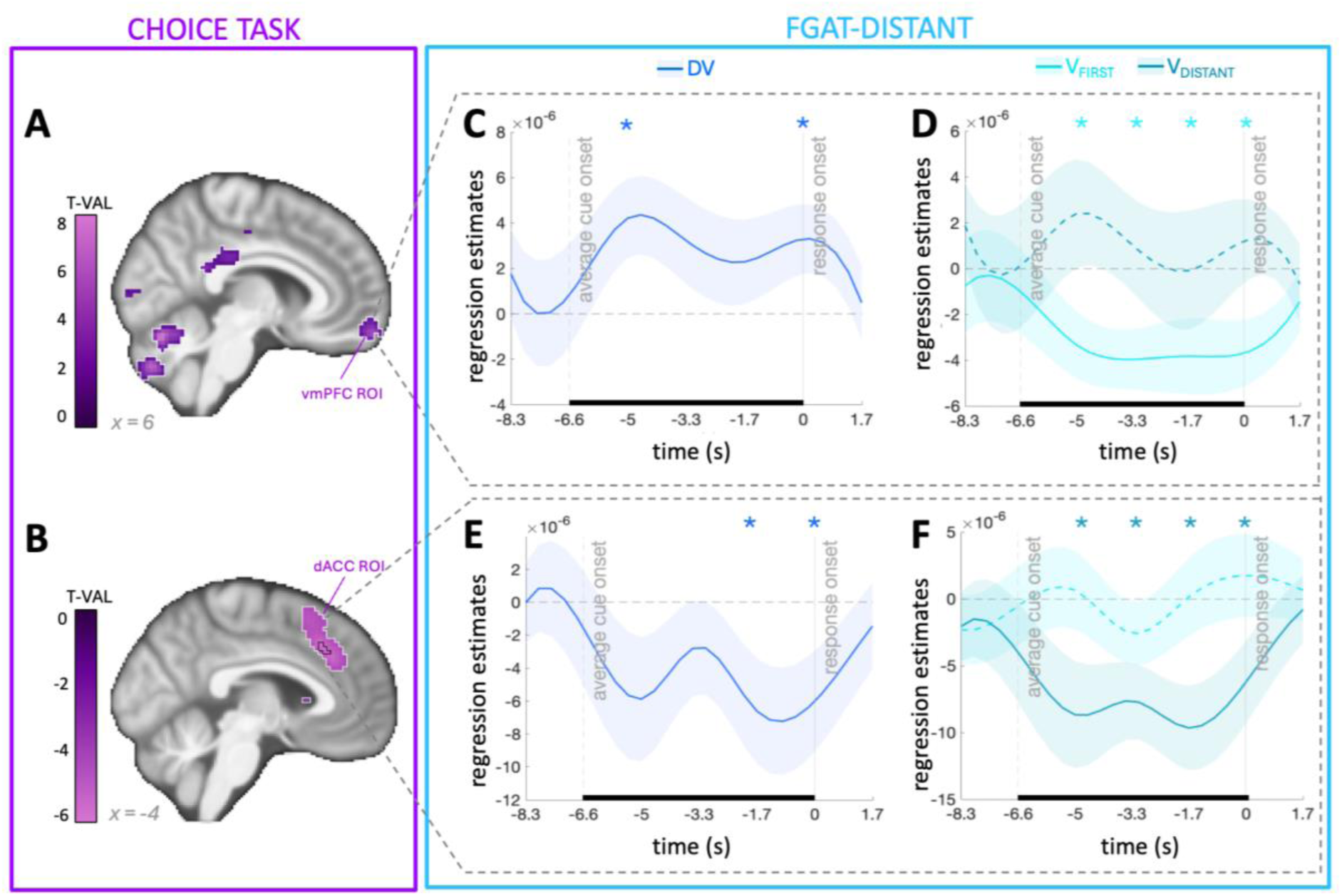
Neural evidence that creative idea production involves idea selection based on the comparison of ideas’ values. **(A & B)** Whole-brain analysis of the **(A)** positive and **(B)** negative neural encoding of the decision value (DV = V_chosen_-V_unchosen_) during the choice task. The color code indicates the T-value of one-sample t-tests, p<0.001 uncorrected. Regions circled in white survived cluster-level FWE correction (p<0.05), the region circled in black survived voxel-level FWE correction (p<0.05). See Table 1 for all significant clusters. The vmPFC and dACC ROIs were used as inclusive masks for the FIR analyses in panels C&D and E&F, respectively. **(C)** Time course of the peri-response regression estimate for the decision value (DV) during the FGAT-distant task, in the vmPFC ROI (shown in panel A) **(D)** Time course of the peri-response regression estimate for the values of FGAT-first and FGAT-distant responses during the FGAT-distant task, in the vmPFC ROI (shown in panel A). **(E)** Time course of the peri-response regression estimate for decision value (DV) during the FGAT-distant task, in the dACC ROI (shown in panel B). **(F)** Time course of the peri-response regression estimate for the values of the FGAT-first and FGAT-distant responses during the FGAT-distant task, in the dACC ROI (shown in panel B). The shaded areas represent the SEM. Solid lines indicate a significant correlation (p<0.05) during the analysis time window (represented here with a horizontal black line), dashed lines indicate non-significant correlation during the analysis time window (p>0.05). The analysis time window is an *a priori* defined time window approximating the mean cue-to-response interval in FGAT-distant trials (RT_FGAT-distant_ = 6.7 ± 0.4 seconds, M±SEM). * indicate significant correlations at specific time points (p<0.05). n=38.

**Table 1:** List of regions encoding decision value during the choice task. Only the results in bold are reported in the text.

| parametric modulation by decision value : positive effects |  |  |  |  |  |  |  |  |
| --- | --- | --- | --- | --- | --- | --- | --- | --- |
| cluster label | cluster size | peak side | x | y | z | Z-score | T-value | cluster p-value |
| cerebellum | 697 | L | -8 | -68 | -22 | 6.37 | 8.67 | <0.001 |
| superior parietal lobule | 247 | L | -28 | -50 | 53 | 4.91 | 5.88 | 0.001 |
| superior parietal lobule | 464 | R | 29 | -45 | 38 | 4.55 | 5.32 | <0.001 |
| temporal occipital fusiform cortex | 608 | L | -38 | -48 | -17 | 4.54 | 5.30 | <0.001 |
| **ventromedial prefrontal cortex** | **165** | **R** | **6** | **62** | **-20** | **4.50** | **5.24** | **0.010** |
| inferior frontal gyrus, pars opercularis | 198 | R | 39 | 15 | 23 | 4.34 | 5.00 | 0.004 |
| posterior cingulate gyrus | 173 | R | 2 | -40 | 20 | 4.20 | 4.79 | 0.008 |
| inferior temporal gyrus | 345 | R | 46 | -48 | -12 | 3.86 | 4.32 | <0.001 |
| middle temporal gyrus | 179 | L | -58 | -10 | -12 | 3.81 | 4.24 | 0.007 |
| Parametric modulation by decision value : negative effects |  |  |  |  |  |  |  |  |
| cluster label | cluster size | peak side | x | y | z | Z-score | T-value | cluster p-value |
| **dorsomedial prefrontal cortex/**<br>**dorsal anterior cingulate cortex** | **586** | **L** | **-4** | **25** | **40** | **05.05** | **6.12** | **<0.001** |
| inferior frontal gyrus, pars opercularis | 137 | L | -48 | 18 | 0 | 4.52 | 5.26 | 0.023 |
| caudate | 166 | R | 9 | 20 | 6 | 4.46 | 5.18 | 0.010 |

Moreover, we found a significant negative effect of DVs in the dACC (Figure 4B): ([−4, 25, 40], one-sample two-tailed t-test: t(37) = 6.12, pFWE < 0.001, see Table 1 for all significant clusters), a region known to negatively encode DVs in decision-making research (Chong et al., 2017; Lopez-Persem et al., 2016; Wunderlich et al., 2009). Given their consistent implication in value-based comparison processes, we hypothesized that the vmPFC and dACC regions would underlie creative idea selection, and used them as regions of interest.

#### 2.4.2. The vmPFC and the dACC activity encode decision values during creative idea production

Next, we used the FGAT-distant task to test whether the brain regions encoding DVs during the choice task also encoded DVs during creative idea production. For these analyses, similarly to the previous RT_FGAT-distant_ analyses, DVs were computed as the difference between the value of the FGAT-distant response (i.e., what was actually chosen during the FGAT-distant) and the value of the FGAT-first response to the same cue word (i.e., a proxy of what was unchosen during the FGAT-distant).

Since we expected the comparison process to occur transiently during FGAT-distant trials, we modeled the fMRI data using a GLM with a finite impulse response (FIR) basis set, allowing us to estimate the event-related time course without imposing a fixed hemodynamic response shape (see Methods). The FIR model estimated the BOLD response at seven time points per trial, spanning 5 TRs (−8.33s) before to 1 TR (+1.66s) after the response press. For statistical analyses, we defined *a priori* a response-locked time window approximating the mean cue-to-response interval in FGAT-distant trials (RT_FGAT-distant_ = 6.7 ± 0.4 seconds, M±SEM), `hereafter referred to as “analysis time window”. Because the FIR time bins are sampled every TR (1.66 s), the closest corresponding window was` −6.64 to 0s relative to the response press, and all statistics were computed within this window for each trial. DV was included as a parametric modulator of the FGAT-distant activity. We applied this analysis within the vmPFC and dACC ROIs using inclusive masks. Note that we performed the same analyses locked on cue onset as control analyses and found that results were mostly non-significant (see Supplementary Results).

First, we found significant positive effects of DVs within the vmPFC ROI at two time points (regression estimate at 5s before response press = 4.10^−6^±2.10^−6^ (M±SEM), one-sample one-tailed t-test: t(37) = 2.40, p = 0.011; regression estimate at response press = 3.10^−6^±2.10^−6^ (M±SEM), one-sample one-tailed t-test: t(37) = 2.13, p = 0.02; Figure 4C, see Table 2 for all results). Since these effects did not survive correction for multiple comparisons (at multiple time points), we investigated whether the effect was on average significant in the analysis time window (i.e., the approximated trial duration, from 6.64s before the response to the onset of the response), and found a significant positive effect of DVs within the vmPFC ROI (analysis time window effect = 3.10^−6^±2.10^−7^ (M±SEM), one-sample one-tailed t-test: t(37) = 1.89 p = 0.03).

**Table 2:**
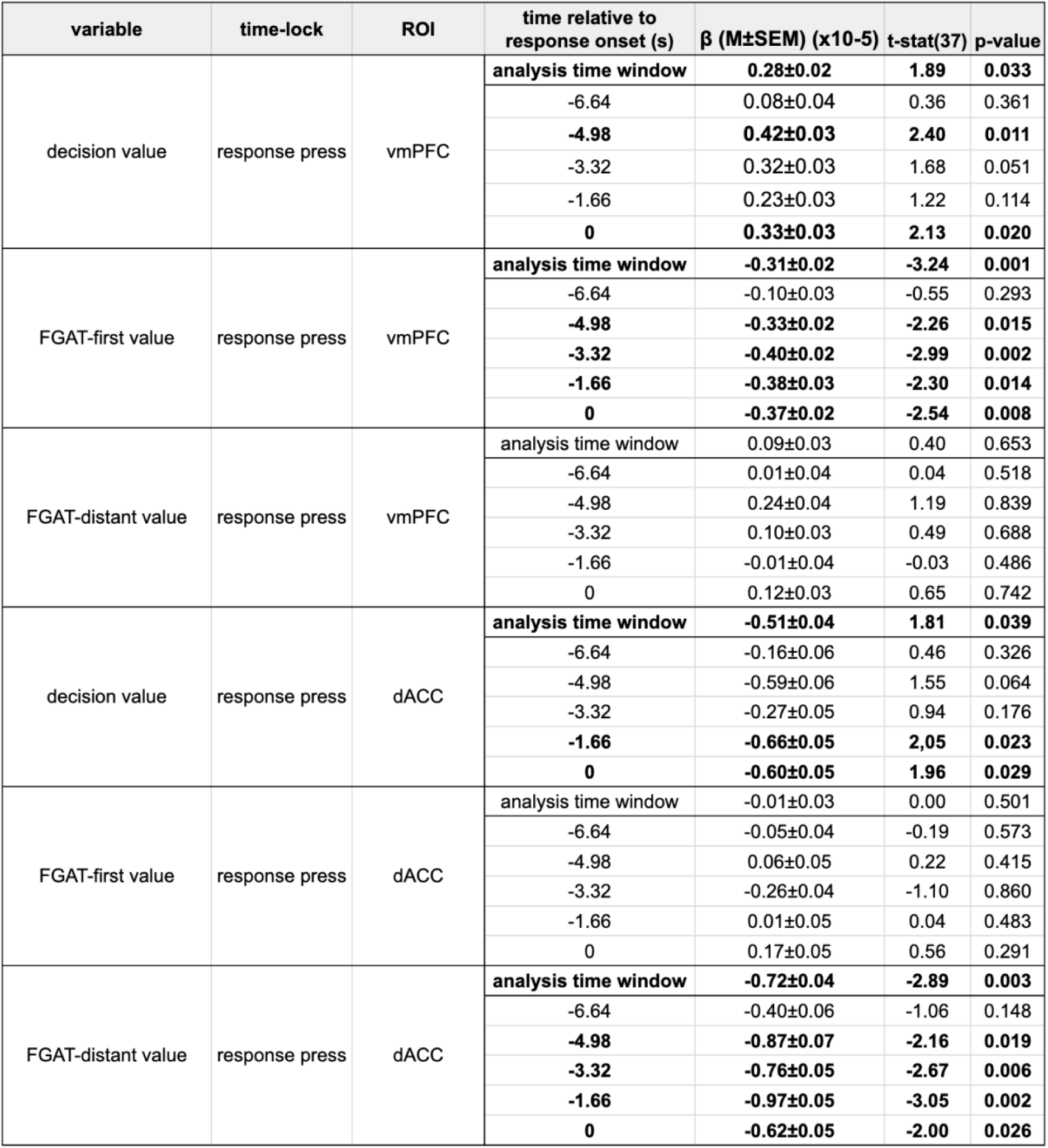
Time-point-wise FIR statistics for parametric modulation in the vmPFC and dACC ROIs during the FGAT-distant, locked on response press. The “analysis time window” is an *a priori* defined time window approximating the mean cue-to-response interval in FGAT-distant trials (RT_FGAT-distant_ = 6.7 ± 0.4 seconds, M±SEM). Results in bold (p<0.05) are significant.

| variable | time-lock | ROI | time relative to response onset (s) | $\beta$ (M±SEM) (x10-5) | t-stat(37) | p-value |
| --- | --- | --- | --- | --- | --- | --- |
| decision value | response press | vmPFC | analysis time window | **0.28±0.02** | **1.89** | **0.033** |
| | | | -6.64 | 0.08±0.04 | 0.36 | 0.361 |
|  |  |  | **-4.98** | **0.42±0.03** | **2.40** | **0.011** |
| | | | -3.32 | 0.32±0.03 | 1.68 | 0.051 |
| | | | -1.66 | 0.23±0.03 | 1.22 | 0.114 |
|  |  |  | **0** | **0.33±0.03** | **2.13** | **0.020** |
| FGAT-first value | response press | vmPFC | analysis time window | **-0.31±0.02** | **-3.24** | **0.001** |
| | | | -6.64 | -0.10±0.03 | -0.55 | 0.293 |
|  |  |  | **-4.98** | **-0.33±0.02** | **-2.26** | **0.015** |
|  |  |  | **-3.32** | **-0.40±0.02** | **-2.99** | **0.002** |
|  |  |  | **-1.66** | **-0.38±0.03** | **-2.30** | **0.014** |
|  |  |  | **0** | **-0.37±0.02** | **-2.54** | **0.008** |
| FGAT-distant value | response press | vmPFC | analysis time window | 0.09±0.03 | 0.40 | 0.653 |
| | | | -6.64 | 0.01±0.04 | 0.04 | 0.518 |
| | | | -4.98 | 0.24±0.04 | 1.19 | 0.839 |
| | | | -3.32 | 0.10±0.03 | 0.49 | 0.688 |
| | | | -1.66 | -0.01±0.04 | -0.03 | 0.486 |
| | | | 0 | 0.12±0.03 | 0.65 | 0.742 |
| decision value | response press | dACC | analysis time window | **-0.51±0.04** | **1.81** | **0.039** |
| | | | -6.64 | -0.16±0.06 | 0.46 | 0.326 |
| | | | -4.98 | -0.59±0.06 | 1.55 | 0.064 |
| | | | -3.32 | -0.27±0.05 | 0.94 | 0.176 |
|  |  |  | **-1.66** | **-0.66±0.05** | **2.05** | **0.023** |
|  |  |  | **0** | **-0.60±0.05** | **1.96** | **0.029** |
| FGAT-first value | response press | dACC | analysis time window | -0.01±0.03 | 0.00 | 0.501 |
| | | | -6.64 | -0.05±0.04 | -0.19 | 0.573 |
| | | | -4.98 | 0.06±0.05 | 0.22 | 0.415 |
| | | | -3.32 | -0.26±0.04 | -1.10 | 0.860 |
| | | | -1.66 | 0.01±0.05 | 0.04 | 0.483 |
| | | | 0 | 0.17±0.05 | 0.56 | 0.291 |
| FGAT-distant value | response press | dACC | analysis time window | **-0.72±0.04** | **-2.89** | **0.003** |
| | | | -6.64 | -0.40±0.06 | -1.06 | 0.148 |
|  |  |  | **-4.98** | **-0.87±0.07** | **-2.16** | **0.019** |
|  |  |  | **-3.32** | **-0.76±0.05** | **-2.67** | **0.006** |
|  |  |  | **-1.66** | **-0.97±0.05** | **-3.05** | **0.002** |
|  |  |  | **0** | **-0.62±0.05** | **-2.00** | **0.026** |

Looking at the encoding of the value of the FGAT-first and FGAT-distant responses individually (Figure 4D), we found that the DV effect was mostly driven by the negative encoding of the value of the FGAT-first response, at all time points from 5s before response press to the onset of response press (see Table 2 for all results). This result was confirmed when testing the effect in the analysis time window (analysis time window effect = −3.10^−6^±2.10^−7^ (M±SEM), one-sample one-tailed t-test: t(37) = −3.2, p = 0.001). Note that there was no significant effect of the value of the FGAT-distant response within the vmPFC ROI (p > 0.05, see Table 2 for more details). These findings confirm our hypothesis that the vmPFC encodes DVs during creative idea production, which is mostly driven by the negative encoding of the value of an unchosen option, the FGAT-first response.

In parallel, we found a significant negative effect of DVs within the dACC ROI at two time points (regression estimate at 1.7s before response press = 6.10^−6^±3.10^−6^ (M±SEM), one-sample one-tailed t-test: t(37) = 2.05, p = 0.023; regression estimate at response press = 6.10^−6^±3.10^−6^ (M±SEM), one-sample one-tailed t-test: t(37) =1.96, p = 0.029; Figure 4E, see Table 2 for all results). The effect was also significant in the analysis time window (analysis time window effect = 5.10^−6^±4.10^−7^ (M±SEM), one-sample one-tailed t-test: t(37) = 1.81, p = 0.04).

Looking at the encoding of the value of the FGAT-first and FGAT-distant responses individually (Figure 4F), we found that the negative encoding of DV was mostly driven by the negative encoding of the value of the FGAT-distant response at all time points from 5s before response press to the onset of response press (see Table 2 for all results). This effect was also significant in the analysis time window (analysis time window effect = −7.10^−6^±4.10^−7^ (M±SEM), one-sample one-tailed t-test: t(37) = −2.89, p = 0.003). Note that the regression estimate for the value of the FGAT-first response was not significant within the dACC ROI (p > 0.05, see Table 2 for more details). These results confirm our hypothesis that the dACC negatively encodes DVs during creative idea production, which is mostly driven by the negative encoding of value of the chosen option, the FGAT-distant response.

Together, these results support the biological validity of our central hypothesis that creative thinking involves the comparison of ideas’ values that drives the selection of the idea with the highest value, thus generalizing classical results of value-based decision-making to the domain of creative thinking. Specifically, this comparison is supported by the positive encoding of DVs in the vmPFC and their negative encoding in the dACC. These results are respectively driven by the negative encoding of the unchosen option in the vmPFC and by the negative encoding of the chosen option in the dACC.

## 3. Discussion

Our framework proposes that during creative thinking, after generating candidate ideas, individuals attribute subjective values to their ideas (valuation), which they then compare (value-based comparison) in order to select and pursue the idea with the highest value (selection). In this study, we aimed to reveal the mechanisms and neural correlates of the value-based comparison of ideas during creative thinking. Based on the decision-making literature, we hypothesized that these neural correlates will involve the BVS and the dACC region. We used a creative idea production task, rating tasks and a choice task to investigate the behavioral and neural mechanisms underlying the value-based comparison of ideas during simple choices between ideas and during creative thinking. After accounting for the fact that the values inferred from the choice task slightly differ from the ones measured via the rating task, behavioral results reveal that response times were influenced by decision values both during simple choices between ideas and during creative thinking, suggesting that a value-based selection process is at play during creative thinking. Next, our neuroimaging analyses identified the neural correlates of the value-based comparison processes. We showed that decision values are encoded positively in the vmPFC and negatively in the dACC both during simple choices and during creative thinking, generalizing typical neural results from the value-based decision-making literature to the context of creative thinking. Overall, our results provide behavioral and neural evidence that idea selection during creative thinking relies on value-based decision-making processes, and bring novel perspectives on the roles of the dACC and vmPFC in creativity.

During the choice task (i.e., when performing value-based comparison), we found that individuals compared values which were slightly but significantly different from those expressed during the rating task. This relates to prior work (Sullivan & Huettel, 2021)on the difference between implicit valuation (here, in the choice task) and explicit valuation (here, in the rating task). Interestingly, it is at odds with previous findings (Gronau et al., 2023; Lopez-Persem et al., 2017) suggesting that valuation parameters driving ratings and choices are similar. Note however that while Lopez-Persem et al. (2017) found that the α valuation parameter was consistent across tasks (in contrast to the current results), the *δ* valuation parameter was higher in the choice task (similar to the current results).

Here, differences in valuation parameters (α and *δ*) indicate that, during choices, individuals put less weight on originality, and showed less preferences for an equilibrium of originality and adequacy over extremes than it would be expected from their ratings. This task effect echoes prior findings suggesting that the importance one places on originality can be context dependent: for example, Long & Pang (2015) found that individuals put more weight on originality when rating answers to open-ended questions, and more weight on adequacy when rating answers to right or wrong questions. Lloyd-Cox et al. (2022) reported that individuals put more weight on originality when rating the creativity of alternative uses for an object, and more weight on adequacy when rating the creativity of social development projects.

Despite their slight difference, both types of valuation parameters correlated with creative performance. This suggests that individuals’ creative performance may rely on either processes, i.e. valuation independent from choices and valuation during choices. Future studies could isolate these processes to determine whether an idea is first evaluated independently before being re-evaluated when compared with alternative ideas, or whether valuation is intrinsic to the comparison process. Regardless, the relationship between either valuation parameters and creative performance supports our hypothesis that individuals’ preferences during idea selection relates to creative performance.

Additionally, we found interindividual differences in the valuation parameters estimated from the choice task, suggesting that people do not all weight originality equally when comparing ideas’ values, which replicates what we previously observed during rating tasks (Battistello et al., 2025; Lopez-Persem et al., 2024; Moreno-Rodriguez et al., 2025). Interestingly, this may underlie the poor selection accuracy reported by several studies (Rietzschel et al., 2006, 2010; Zhu et al., 2017, 2021), as interindividual differences in weightings of originality could explain the gap between participants’ and experts’ selection.

Next, we searched for a signal of a value-based comparison process during choices and creative idea production. In the choice task, participants responded more quickly when decision values were larger: stronger preferences led to faster, i.e. easier, choices. This reflects classical value-based decision-making findings, where choice task response times systematically decrease as preference strength increases (De Martino et al., 2013; Konovalov & Krajbich, 2019). Finding this pattern during a creative choice task confirms that a value-based comparison process is at play during creative idea selection. Importantly, we found the same pattern during creative idea production, i.e., when individuals were producing an idea (FGAT-distant) that outcompeted a prior response (FGAT-first). This second result suggests that the value-based comparison process is not restricted to choice tasks, but is also at play during creative idea production. Note that, in previous studies from the lab (Battistello et al., 2025; Lopez-Persem et al., 2024; Moreno-Rodriguez et al., 2025), we showed that individuals produced the ideas they preferred faster, and interpreted that result as a general energisation effect. The current results provide more information about this energization effect, suggesting that the acceleration of RT might result from an easier comparison process, reflecting the strength of preferences, and energizing behavior.

In addition to this behavioral evidence, neuroimaging results revealed some brain correlates of the value-based comparison process. In particular, decision values were encoded positively in the vmPFC and negatively in the dACC. This extends to creativity neuroscience the neural findings of classical decision making research (Grueschow et al., 2015; Hare et al., 2011; Hunt et al., 2014; Kolling et al., 2012; Plassmann et al., 2007; Rushworth et al., 2012; Shenhav et al., 2013), which usually demonstrate the encoding of decision values of economic or dietary decisions in these regions. These results are in line with the common neural currency theory, which states that the values of different and disparate types of items (i.e, food, nature and here, creative items) can be represented in the same brain regions and allows for their comparison (Levy & Glimcher, 2012).

Regarding the dACC, note that this region lies at the intersection of two networks repeatedly implicated in creativity: the Executive Control Network (ECN) and the Salience Network (SN) (Yeo et al., 2011). Observing dACC activity during the production of creative ideas confirms the involvement of these networks in creativity but also challenges, or expands, their traditionally proposed roles. In the neuroscience of creativity, the ECN is usually associated with idea evaluation and the SN is usually associated with regulating the balance between ECN and DMN activity (Beaty et al., 2015, 2016), yet here dACC supports idea comparison. Further research will be needed to clarify to which network the dACC region is connected to, intrinsically and during creative idea selection, and whether the SN (or ECN) is involved in value-based comparison or in another creativity process.

Interestingly, during the FGAT-distant, the negative encoding of the decision values in the dACC was mostly driven by the negative encoding of the value of the FGAT-distant option. This negative encoding of the chosen option’s value in the dACC during a choice is consistent with previous results suggesting that the chosen value is represented negatively in this brain region (Lopez-Persem et al., 2016; Wunderlich et al., 2009), akin to an “engage value” (Kolling et al., 2016), or reflecting the difficulty of the decision at play (Gloy et al., 2020; Shenhav et al., 2013, 2013; Volz et al., 2005). We might have also expected the positive encoding of the FGAT-first (unchosen) response, given that this region is known to encode alternative or search value (Kolling et al., 2016). One possible explanation for its absence in our results is that the value for first response that comes to mind may not represent genuine alternative or search value (Rushworth et al., 2012). Future studies using a fluent version of the FGAT (where participants report and rate all of the ideas that come to their mind) will be better suited to track how this brain region encodes the values of alternative options considered by individuals during creative thinking.

In contrast, during the FGAT-distant, the positive encoding of the decision values in the vmPFC was mostly driven by the negative encoding of the value of the FGAT-first option. Although this negative encoding echoes previous results showing that the unchosen value is reflected negatively in the vmPFC (Boorman et al., 2009, 2013), not observing a significant positive encoding of the FGAT-distant response value was surprising, for two main reasons. First, chosen value encoding is usually observed in choices (Kolling et al., 2012; Wunderlich et al., 2010), and second, this region was observed when using likeability of the FGAT-distant response in our previous study using the same fMRI dataset (Moreno-Rodriguez et al., 2025). However, the vmPFC ROI in the current study is more anterior than the vmPFC ROI of our previous study. Moreover, the timing of the analysis is different: here, the analysis time window is locked on the response, while our previous study used the whole thinking time period. Furthermore, our two studies differ in the way values were computed: the current study uses values computed from originality and adequacy ratings as parametric modulator, while our previous study’s values were likeability ratings. Regardless, the negative encoding of FGAT-first response value provides evidence that a rejected alternative response value is encoded during creative thinking, suggesting that it was considered by participants when having to select a response.

Overall, finding that decision values were encoded during FGAT-distant brings forth neuroimaging evidence in line with our hypothesis that creative idea production relies on a value-based comparison process and identifies brain regions supporting this process.

### Limitations and future directions

One limitation of the present study is that, while it demonstrates the neural encoding of decision value during creative idea production, thus suggesting that value-based comparison is at play, it does not establish that this process is necessary for idea production. Moreover, the observed behavioral (response times) and neural signals (dACC activation) may reflect difficulty or confidence rather than the encoding of decision value specifically, which complicates the interpretation of our findings. Nevertheless, since decision value, difficulty and confidence all implicate decision-making processes, this interpretative ambiguity does not undermine the core conclusion that value-based comparison mechanisms operate during creative thinking. Furthermore, although we show that individual differences in valuation parameters (α_choice_) relate to creative performance, the results are correlational and do not allow us to infer a causal relationship. For example, individuals with a higher weighting of originality may also differ in other traits (such as motivation, or cognitive flexibility) that might contribute to their creative abilities. In other words, we cannot yet tell whether valuing originality leads to higher creative performance, or if it is simply a characteristic of already more creative individuals. Overall, while the present study shows that value-based decision-making processes are *featured* during creative idea production, it does not establish whether these processes are *necessary* for creativity. Future studies could build on our value-based framework and directly test the mechanistic role of value-based processes in creativity, for example by modulating valuation parameters and observing the impact on creative performance in a behavioral intervention.

### Conclusion

Together, our findings provide converging behavioral and neural evidence that creative idea production is driven by value-based decision-making mechanisms. By combining a creative thinking task, rating tasks and a choice task, our results not only extend classical decision-making findings to the field of creativity, but also reveal that the selection of ideas during creative thinking results from a value-based comparison process, supported by core value-based decision-making brain circuits. Finally, our results suggest that giving more importance to originality when comparing and selecting ideas may be a key driver of creative performance. This study paves the way to investigating whether the modulation of these valuation parameters can directly boost creative performance, which will be necessary to establish the causal role of value-based processes in creativity.

## 4. Methods

### 4.1. Participants

The present study uses the behavioral and fMRI data from two previous studies (Lopez-Persem et al, 2024, Moreno-Rodriguez et al, 2025). An official ethics committee approved both these studies (CPP Ouest II – Angers). We recruited and tested 71 participants at the PRISME platform (subgroup A) and 40 participants at the CENIR platform (subgroup B) of the Paris Brain Institute (ICM). All participants were French native speakers, right-handed, with correct or corrected vision, and no neurological or psychological disease history. They gave informed consent and were compensated for their participation. We excluded from all analyses two participants of subgroup A (because of a misunderstanding of the instructions) and one participant of subgroup B (because we interrupted MRI scanning after they suffered from a claustrophobic episode). We excluded another participant of subgroup B from MRI analyses (because of a technical issue during MRI scanning, that did not impact the behavioral data). 69 subjects of subgroup A (28 males and 41 females, age = 25.8±0.5 (M±SEM), years of education = 17.0±0.2 (M±SEM)), and 39 subjects of subgroup B (20 males and 19 females, age = 26.5±0.6 (M±SEM), years of education = 16.8±0.4 (M±SEM)) were included in the analyses. The final total sample comprised 108 participants (48 males and 60 females, age = 26.0±0.4 (M±SEM), years of education = 16.9±0.2 (M±SEM)). Participants of subgroup A were compensated with 40€ for a four-hour testing session that included cognitive and creativity tests outside the scanner. Participants of subgroup B were compensated with 110€ for a four-hour testing session that included MRI scanning and cognitive and creativity tests inside and outside the scanner.

### 4.2. Experimental design

Participants of subgroup A completed an FGAT-first, FGAT-distant, a likeability rating task, a choice task, an originality and adequacy rating task and a battery of creativity tests outside the MRI. Participants of subgroup B completed an MRI session composed of an anatomical T1 scan, four task-based fMRI scans (while performing the FGAT-first, FGAT-distant, the likeability rating task and the choice task) and a resting-state fMRI scan. Participants of subgroup B then completed the originality and adequacy rating task and the battery of creativity tests outside the MRI. We used Matlab (MATLAB. (2020). 9.9.0.1495850 (R2020b). Natick, 624 Massachusetts: The MathWorks Inc.) to program the tasks and the Qualtrics software (Qualtrics, Provo, UT, USA. https://www.qualtrics.com) to implement the battery of creativity tests.

#### 4.2.1 Free Generation of Associates Task

The Free Generation of Associates Task (FGAT, Figure 1B) is a remote thinking task reported to capture critical aspects of creativity (Bendetowicz et al., 2018; Prabhakaran et al., 2014, Lopez-Persem et al., 2024, Moreno-Rodriguez et al., 2025, Battistello et al., 2025). In our experimental design, the task comprised two successive conditions, each comprising 5 training trials and 62 randomized test trials. At each trial, a cue word was displayed. In the “first” condition, participants had up to 10 seconds to give the first word that came to mind after reading the cue word (for example, participants might answer “father” to the cue word “mother”). In the “distant” condition, they had up to 20 seconds to provide a distant word, so the cue and response words would result in a creative word association (for example, participants might answer “nature” to the cue word “mother”). Outside the MRI (subgroup A), participants pressed the spacebar of the keyboard when they had an answer in mind, then they typed their answer and validated it. In the MRI (subgroup B), each trial started with a fixation cross for 1.6 to 3.2 seconds, jittered with a uniform distribution, and participants pressed the response button with their right index finger when they had an answer in mind. Because participants could not type their response while in the MRI, they said the word out loud into a microphone. The experimenter typed the response and displayed it on the participant’s screen. Participants then had the opportunity to repeat their response in case the experimenter had misheard (by pressing the response button again with their right index finger) or to validate the response (by pressing the validation button with their right ring finger). The exact instructions given to the participants are in the Supplementary Methods.

#### 4.2.2. Rating tasks

We split the rating task in two: a likeability rating task (performed inside the scanner by subgroup B) and an originality and adequacy rating task (performed outside the scanner by all participants). Each rating task counted five training trials and a variable number of test trials (M±SEM = 145±2), depending on the validity of the participant’s FGAT answers (see Supplementary Methods for more details). For both rating tasks, the trials were composed of the following: 66% of the trials were evenly made up of the participant’s valid FGAT-first and FGAT-distant associations; 27% of the trials were a sample of frequent and rare responses taken from another study using the same FGAT cue words; and finally 7% of the trials were responses that were totally unrelated to the cue words they were associated with (for example: ‘cow’ and ‘inverse’). We added these associations to cover a wide range of adequacy and originality ratings, ensuring a robust estimation of likeability with sufficient statistical power.

The first rating task, i.e. the likeability rating task (Figure 1C), was performed in the MRI scanner by subgroup B, and outside the MRI by subgroup A. Participants had to indicate how much they liked the association or how satisfying it was in the context of the FGAT-distant condition. First, a cue word (for example, “mother”) was displayed alone for 0.3 seconds (outside the MRI) or for 0 to 0.5 seconds (in the MRI: all display times varied between trials and were determined by a random jitter with a uniform distribution). Then, a response appeared under the cue word. This association (for example, “mother-nature”) remained alone for 0.4 seconds (outside the MRI) or between 1.6 to 3.2 seconds (inside the MRI). Then, a rating scale appeared below the association. The rating scale’s low to high values were represented from left to right, with the words “not at all” and “very much” located at the extremities but without any numerical values. A segment indicated the middle of the scale, which was divided into 101 hidden steps, later converted into ratings ranging between 0 and 100. We displayed a heart icon (for likeability) under the rating scale. Outside the MRI, participants provided a rating using the left and right arrows of their keyboard, and validated their answer by pressing the spacebar. In the MRI, participants had to move a cursor by pressing the first and second MRI response buttons with the right index and right middle finger and then validate the rating by pressing the third button with their right ring finger. Once the response was validated, the subsequent trial began. In the MRI (subgroup B), each trial started with a fixation cross displayed for 1.6 to 3.2 seconds. The exact instructions given to the participants are in the Supplementary Methods.

The second rating task, i.e. the originality and adequacy rating task (Figure 1D), was completed after the choice task and outside the MRI for all participants. Again, each trial displayed a two-word association (for example, “mother-nature”) and asked participants to subsequently rate how original and how adequate it was. The order of originality and adequacy ratings was counterbalanced across trials. Each trial started with the screen displaying a cue word. After 0.3 seconds, a response word appeared under the cue, and after 0.4 seconds, a scale appeared under the two-word association. The scale was similar to the likeability rating scale except for the icon displayed underneath: a star for originality and a target and arrow for adequacy. Participants rated the association by moving the slider across the scale using the left and right arrows of the keyboard and validated by pressing the spacebar. Then, a new scale appeared underneath the same association, indicating the second dimension (adequacy or originality). Participants provided their ratings similarly and then started the subsequent trial.

We told participants to use the whole scale throughout the task. No time limit was applied, but the instruction was to respond spontaneously. The exact instructions given to the participants are in the Supplementary Methods.

#### 4.2.3. Choice task

The choice task was performed in the MRI by subgroup B, and outside the MRI by subgroup A. It comprised five training trials and an average of 187 randomized test trials (the exact number varied between participants depending on the variability of their FGAT answers and ratings). Each trial displayed two possible responses to an FGAT cue, with the cue word on the top of the screen and the two response options words below, on the same horizontal line (for example, “mother” was displayed on top of “nature” and “father”). Note that some trials exhibited two associations given by the participant (i.e. their FGAT-first versus their FGAT-distant answer for the same cue), but some other trials opposed their own association to an association taken from another study from the lab using the same FGAT cue words, and some other trials opposed two associations from that database. Participants had to choose which of the two responses they would have preferred to provide for a given cue in the FGAT-distant condition. Outside the MRI, participants from subgroup A chose the left or the right response using the left or right arrow of their keyboard. In the MRI, participants from subgroup B pressed the first or second MRI response buttons with the right index or right middle finger. Choices were saved as soon as a key/button was pressed and did not require validation. No time limit was applied, but the instruction was to respond spontaneously. The exact instructions given to the participants are in the Supplementary Methods.

#### 4.2.5. Battery of creativity tests

The battery comprised a daily-life creativity questionnaire, self-reports, and established creativity tasks belonging to three main frameworks of creativity - the divergent thinking approach, the associative theory, and the insight problem-solving approach.

##### Inventory of Creative Activities and Achievements (ICAA)

the inventory of creative activities and achievements was first designed by Diedrich et al. (2018) as an ecological measure of creativity. It focuses on eight domains of creativity (literature, music, crafts, cooking, sports, visual arts, performing arts, and science and engineering). For each domain, participants reported their level and their frequency of engagement over the past ten years. They also specified their level of creative accomplishment in each of these domains. We computed the ICAA score as the mean of the creative activities score and the creative achievements score. First, for the creative activities score, we summed the frequency at which the participant had engaged in each of the eight creativity domains using an ordinal scale ranging from 0 (never) to 4 (more than ten times). Then, for the creative achievements score, we summed the participants’ levels in each of the eight creativity domains. These ranged from 0 (“never engaged in this domain”) to 10 (“I have already sold some of my work in this domain”). Finally, we averaged the two scores into a unique ICAA score per participant.

##### Drawing task

the drawing task (Barbot, 2018) comprised a training trial and 12 test trials. We instructed participants to include incomplete shapes in creative drawings. The 12 incomplete shapes were relatively similar: they were all composed of 4 lines, either straight, curved or right-angled, and exhibited symmetry (cf. (Barbot, 2018) for an example). Participants completed the 12 drawings at their own pace. Then, they reviewed each drawing and gave each one a verbal title. We followed the consensual assessment technique that is advised for scoring such a creativity task (Ceh et al., 2022). We designated four judges from the lab who were familiar with creativity judgments but had not seen the data before the rating. For each of the twelve shapes, the judges first saw the incomplete shape, then each participant’s drawing using that shape, in a blind and randomized order. Then, the judges gave each drawing a rating between 0 (not creative at all) and 4 (extremely creative). We tested the inter-judge independent rating consistency using an intraclass correlation analysis, which gives the mean correlation between the judges’ ratings (r = 0.8239 ± 0.02; M±SEM).

##### Combination of Associates Task

Bendetowicz et al. (2017) first developed the Combination of Associates Task (CAT) by adapting the remote associative task from Mednick (1962). After five training trials, participants performed 40 randomized test trials, each displaying three cue words with no apparent association: they had to find a fourth word that linked all of them (for example, the link between “map”, “dig”, and “discover” is “treasure”). The CAT score used for the analyses was the participants’ total number of accurate answers.

##### Associative fluency task

in this task, adapted from Benedek et al. (2012), participants had two minutes to type as many words related to a cue word as possible. We chose this associative design over simple category or letter fluency tasks to measure creative abilities on top of fluency. We selected six cue words (“garden”, “wine”, “rock”, “opinion”, “call”, and “finger”) among the FGAT cues for their diversity in steepness. The steepness is the ratio of the frequency of the most frequent answer over the frequency of the second most frequent answer: a steep cue word (e.g., “cat”) has a very common first associate (“dog”); a flat cue word has several equally-common first associates. We measured typing speed and accounted for it in the fluency analysis. The score used for the analyses was the total number of answers given to a cue word, averaged across all cue words.

##### Alternative uses task (AUT)

the alternative uses task used in this study derives from Guilford’s (1967) work and is a commonly used test of divergent thinking. For each of three everyday items (a tire, a knife, and a bottle), participants had 3 minutes to list as many alternative uses of this item as possible. Then, they chose the three most creative and their three favorite answers, as previous research reported that using the top 3 scorings effectively assesses creativity (Benedek et al., 2013).

### 4.3. Behavioral analyses

#### 4.3.1. Model comparison, parameters estimates and correlations

##### Model space definition

To identify which decision values were computed during the choice task, we compared four versions of a choice model. These four models were all based on a softmax function.

The softmax function explains the probability that a participant will choose an option in function of the decision value (DV). DV is defined as the difference between the values of the two choice options (DV = V _option 1_ - V _option 2_). Here, we looked at the probability of choosing the FGAT-distant option in choices opposing FGAT-first and FGAT-distant responses (Equation A), thus DV = V_FGAT-distant_ - V_FGAT-first_. Note that these responses could be either the participant’s or responses taken from an external database (Altmayer et al., 2025). In its simpler form, the softmax function includes one parameter: the inverse temperature (*β* parameter, the expected stochasticity, or incoherence, in individuals’ choices).

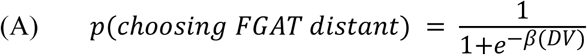

While the four models relied on this softmax function, they differed in two ways. First, they differed in the computation of the values used in their DVs. All four value computations were based on the following Constant Elasticity of Substitution (CES) function (Equation B).

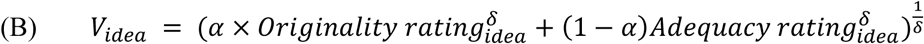

Previous papers from the lab (Lopez-Persem et al., 2024, Battistello et al, 2025) used model comparison and demonstrated that it was this function of originality and adequacy that best fitted participants’ likeability ratings. In this function *⍺* captures the weight given to originality relative to adequacy in likeability ratings. If *⍺* is greater than 0.5, likeability is more strongly driven by originality than adequacy, meaning the participant prioritizes originality (and vice versa if *⍺* is below 0.5). *δ* captures the preference for a balance between originality and adequacy. If *δ* is lower than 1, likeability increases when both dimensions are high, indicating a preference for equilibrium. If *δ* is greater than 1, likeability increases when one dimension is much higher than the other, reflecting a preference for extremes.

If all models used this CES function, models 1 and 2 used fixed parameters to compute values: *⍺*_rating_ and *δ*_rating_ previously estimated from the likeability ratings. Models 3 and 4, on the other hand, used free parameters, i.e., parameters to be directly estimated from the choices: *⍺*_choice_ and *δ*_choice_.

Second, the four models differed in whether they included a free parameter *γ*, which captures whether the participants exhibit an additive bias towards FGAT-distant ideas and against FGAT-first ideas (*γ* > 0) or against FGAT-distant ideas and towards FGAT-first ideas (*γ* < 0). Models 2 and 4’s softmax function featured this parameter, while models 1 and 3’s did not. In summary, the models were defined as follows:

***Value-before-choice models:***

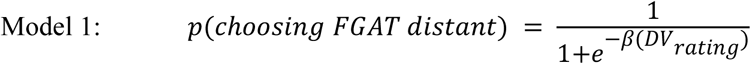

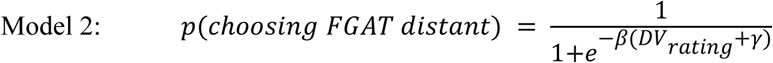

***Value-during-choice models:***

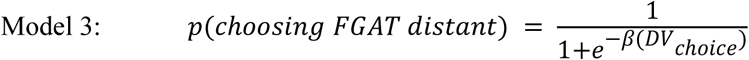

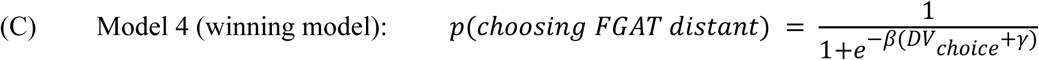

In these equations, DV = V_FGAT-distant_ - V_FGAT-first_, where the values V are computed as shown in Equation B. DV_rating_ uses V computed with fixed parameters *⍺*_rating_ and *δ*_rating_, while DV_choice_ uses V computed with free parameters *⍺*_choice_ and *δ*_choice_. The inverse temperature parameter (*β*) was defined as a free parameter in all models.

##### Model comparison

We conducted a model comparison using the Variational Bayesian Analysis toolbox for MATLAB (https://mbb-team.github.io/VBA-toolbox/), which applies a variational Bayesian framework based on the Laplace approximation (Daunizeau et al., 2009; Stephan et al., 2009). This approach iteratively computes a free energy bound on the model evidence, balancing model fit against complexity in a principled manner (Friston et al., 2007; Penny, 2012). It also yields posterior distributions over the model parameters, assuming initial Gaussian priors. At the group level, individual log-model evidence values were submitted to a random-effects Bayesian model selection procedure (Rigoux et al., 2014; Stephan et al., 2009). This analysis outputs both the exceedance probability (Xp), which quantifies the likelihood that a given model is more frequent than any other in the population, and the estimated model frequency (Ef), which reflects the expected proportion of participants best described by each model. A model is typically considered the winning model if it has both the highest exceedance probability and an Xp above a threshold (here, 0.95), indicating strong evidence for its predominance in the population. Balanced accuracy range provides an absolute measure of the fit of the model across the model space, with values above 0.5 reflecting its ability to predict behavioral responses above chance level.

##### Analyses of the estimated parameters

After identifying the winning model, we investgated the *⍺*_choice_, *δ*_choice_ and *γ* parameters estimated for each participant. Using one-sample, two-tailed t-tests, we tested each parameter against their threshold (0.5 for *⍺*_choice_, 1 for *δ*_choice_ and 0 for *γ*). Then, we compared the *⍺*_choice_ and *δ*_choice_ parameters to the *⍺*_rating_ and *δ*_rating_ parameters using paired two-tailed t-tests. We also tested the Pearson correlation between *⍺*_choice_ and the *γ* parameter. Finally, we tested the correlation between *⍺*_choice_ and *⍺*_rating_ and between *δ*_choice_ and *δ*_rating_.

#### 4.3.2. Relationship between decision values and response times

We performed two linear regressions of RTs against DVs, first in the choice task, then in the FGAT-distant.

In the first linear regression, DVs were defined for each choice trial as the difference between the value of the association that was chosen, and the value of the association that was not chosen by the participant. These values were computed using the CES function (Equation B), *⍺*_choice_, *δ*_choice_ and the participant’s originality and adequacy ratings of these associations. Importantly, we only included the trials that opposed two associations given by the participant, excluding the trial where at least one of the choice options was an association taken from the database of another study from the lab (cf. choice task methods). Response times were defined as the time between cue display and response button press. We excluded choice trials with response times shorter than 0.3 seconds which are likely to reflect errors rather than actual value-based deliberation (0.5 % of trials).

In the second linear regression, DVs were defined for each FGAT-distant trial as the difference between the value of the “chosen answer” (the FGAT-distant response of this trial) and the value of the “unchosen answer”. As a proxy for this, we used the FGAT-first response given to this trial’s cue word in the FGAT-first condition. Response times were defined as the time between cue display and response button press. We excluded FGAT-distant trials with response times shorter than 0.3 seconds which are likely to reflect errors (2% of trials).

Then, for each linear regression analysis, we z-scored DVs and fitted the linear regression at the participant level using MATLAB’s fitlm function. Then, we tested the resulting regression coefficients for significance at the group level using one-sample two-tailed t-tests. For figures 2B and 2C, we binned each participant’s DVs into 10 equally spaced intervals ranging from −100 to 100 (DV extreme values). For each bin, we calculated the mean DV and the mean response time, and plotted these averages.

### 4.4. Neural analyses

#### 4.4.1. MRI data acquisition and preprocessing

##### Scanning parameters

We acquired neuroimaging data on a 3T MRI scanner (Siemens 3T Magnetom Prisma Fit) with a 64-channel head coil.

Four functional runs were acquired (FGAT-first, FGAT-distant, likeability rating task, choice task) using multi-echo echo-planar imaging (EPI) sequences. The number of whole-brain volumes per run varied between tasks but also between participants since all tasks were self-paced (mean, [min;max]: FGAT-first = 379, [317;454], FGAT-distant = 612, [359;856]; likeability rating task = 800, [650;1012] and choice task = 662, [525;821]). We did not record any dummy scans and, therefore, did not discard any volumes. The functional runs used the following parameters: repetition time (TR) = 1660 ms; echo times (TE) for echo 1 = 14.2 ms, echo 2 = 35.39 ms, and echo 3 = 56.58 ms; flip angle = 74°; 60 slices, slice thickness = 2.50 mm; isotropic voxel size of 2.5 mm; Ipat acceleration factor = 2; multiband = 3; and interleaved slice ordering.

After the EPI acquisitions, we acquired a T1-weighted structural image with the following parameters: TR = 2300 ms, TE = 2.76 ms, flip angle = 9°, 192 sagittal slices with a 1-mm thickness, isotropic voxel size of 1 mm, Ipat acceleration factor = 2.

Finally, we added a resting-state fMRI session of 10 min (360 volumes) with the same acquisition parameters as the task runs.

##### Preprocessing

We performed the preprocessing of the on-task fMRI data separately for each run and the resting-state data using the afni_proc.py pipeline from the Analysis of Functional Neuroimages software (AFNI; https://afni.nimh.nih.gov). The different preprocessing steps of the data included slice timing correction and realignment to the first volume (computed on the first echo). We then combined the preprocessed data using the TE-dependent analysis of multi-echo fMRI data (TEDANA; https://tedana.readthedocs.io/), version 0.0.9a1 (DuPre et al., 2021, Kundu et al., 2013). We co-registered the resulting data on the T1-weighted structural image using the Statistical Parametric Mapping (SPM) 12 package running in MATLAB (MATLAB R2017b, The MathWorks Inc., USA). We normalized the data to the Montreal Neurological Institute template brain using the transformation matrix computed from the normalization of the T1-weighted structural image with the default settings of the computational anatomy toolbox (CAT 12; http://dbm.neuro.uni-jena.de/cat/) (Gaser et al., 2024) implemented in SPM 12. We entered the resulting normalized data from the task-based fMRI in generalized linear models (GLMs) in SPM. In this analysis, we entered 24 motion parameters and 42 physiological noise parameters as confounds regressed from the blood-oxygen-level-dependent (BOLD) signal. In this preprocessing analysis, we used the Matlab PhysIO Toolbox (Kasper et al., 2017) (version 5.1.2, open-source code available as part of the TAPAS software collection: https://www.translationalneuromodeling.org/tapas, Frässle et al., 2021) to generate nuisance regressors for the GLM. The regressors were composed of (i) physiological (cardiac pulse and respiration) recordings used to generate RETROICOR (Glover et al., 2000) regressors, (ii) White Matter (WM) and CerebroSpinal Fluid (CSF) masks from the anatomical segmentation used to extract components from compartments of non-interest using Principal Component Analysis (PCA) and (iii) motion parameters, composed of standard motion parameters, first temporal derivatives, standard motion parameters squared, and first temporal derivatives squared. Outlier volumes were eliminated when the Framewise Displacement (FD) exceeded 0.5mm. Then, we used GLMs to explain preprocessed time-series at the individual level.

#### 4.4.2. Statistical analyses of BOLD responses in choices and FGAT-distant

To examine the neural correlates of value-based selection during choices and during FGAT-distant, we used a general linear model (GLM) to explain pre-processed timeseries at the individual level. The first model (GLM1) included a boxcar function modelling the signal between the cue display and the button press in the choice task, which was parametrically modulated by the DV (difference between the value of the chosen response and the value of the unchosen response). These values were computed using the CES function (Equation B), *⍺*_choice_, *δ*_choice_ and the participant’s originality and adequacy ratings of these associations. The regressor was convolved with a canonical hemodynamic response function (HRF). Regression coefficients were estimated at the individual level and then taken to group-level random-effect analysis using one-sample two-tailed t-test.

The second model, GLM2, was similar except that it was used to model timeseries from the FGAT-distant condition. Because the exact timing of the decision process was less constrained in this task, we used a finite impulse response function (FIR) approach rather than assuming a canonical HRF. The FIR model estimated the BOLD response at seven time points per trial, from 5 TRs (−8.33s) before to 1 TR (1.66s) after the response press. Mirroring the choice task, DVs were entered as a parametric modulator and were defined as the difference between the value of the response produced in the FGAT-distant condition (the chosen option) and the value of the response produced in the FGAT-first condition (a proxy for an unchosen option). These values were computed using the CES function (Equation B), *⍺*_choice_, *δ*_choice_ and the participant’s originality and adequacy ratings of these associations.

GLM3 and GLM4 were exactly as GLM2 except that instead of using DV, only the value of the FGAT-first response (GLM3) and FGAT-distant response (GLM4) respectively, were used as parametric modulators.

GLM1 allowed us to define the ROIs used in GLM2, GLM3 and GLM4, with a threshold of p < 0.05, controlling for family-wise errors (FWE) at the cluster level (using cluster thresholds of 120 for positive contrast and 137 for negative contrast in GLM1). For GLM2, GLM3 and GLM4, regression coefficients of each time point were extracted within the ROI defined in the choice task.

We computed the mean and standard error of the mean (shaded areas in Figure 3) of each point’s regression estimates. Then, we tested the significance of each data point’s regression estimate using one-sample, one-tailed t-tests. Next, since the previous multiple correlations were uncorrected, we also tested the significance of the regression across a time window, which was a priori defined as the average duration of FGAT-distant trials across participants (average time between the cue onset and the button press signaling that the participant had found a response to the cue = 6.7 ± 0.4 s, which amounts to 4 TRs before response press), using one-sample, one-tailed t-tests.

Note that not all 62 FGAT-distant trials were included in these FIR analyses: depending on the participant, the number of included trials ranged between 34 and 59, as some associations had not been rated after controlling for FGAT-first and FGAT-distant similarity (see Supplementary Methods for more details).

### 4.5. The relationship between creativity and the parameters estimated using the choice task

#### 4.5.1. The relationship between creative performance and the parameters estimated using the choice task

We investigated the relationship between the parameters estimated using the choice task (*⍺*_choice_, *δ*_choice_ and the *γ* parameter) and creative performance by looking at the parameters’ Pearson correlation with the number of unique answers in FGAT-distant.

#### 4.5.2. The relationship between creative abilities and the parameters estimated using the choice task

Then, we investigated the relationship between the parameters estimated using the choice task (*⍺*_choice_, *δ*_choice_ and the *γ* parameter) and creative abilities by looking at the parameters’ correlation with the scores in the battery of creativity tests. We decreased the dimensionality of these scores using a principal components analysis (PCA).

##### PCA

We performed a PCA on the data from a drawing task, an Inventory of Creative Activities and Achievements (ICAA), an Alternative Uses Task (AUT), a Combination of Associates Task (CAT) and an associative fluency task. We conducted a Bartlett’s test and a Kaiser-Meyer-Olkin (KMO) measure of sampling adequacy, to test the suitability of the data for a PCA. We tested the statistical significance of the χ^2^ parameter using a one-sample, two-tailed t-test. We also tested the general KMO measure for suitability (Table S1). Finally, we generated a scree plot which showed a clear elbow suggesting that two components were sufficient to capture most of the variance, which, together, accounted for 60.5% of the variance. The first principal component (PC1) explained 37.5% of the variance, and the second principal component (PC2) explained 23% of the variance (Table S1).

PC1 was primarily associated with high positive loadings on the ICAA and the associative fluency, followed by the drawing task, indicating that PC1 captures divergent thinking abilities and real-life creative abilities. PC2, on the other hand, was primarily associated with high positive loadings on the CAT, the drawing task and the AUT, indicating that it captures both convergent and divergent abilities.

##### Correlation analyses

Then, we tested the Pearson correlation between the parameters estimated using the choice task (*⍺*_choice_, *δ*_choice_ and the *γ* parameter) and each of the two identified PCA components (Table S2 and Figure 1E-G).

## Notes

### Competing Interest Statement

The authors have declared no competing interest.

## References

Altmayer, V., Ovando-Tellez, M., Bieth, T., Batrancourt, B., Rametti-Lacroux, A., Bernardaud, L., Moreno-Rodriguez, S., Vigreux, L., Ledu, V., Garcin, B., Migliaccio, R., Le Ber, I., Lopez-Persem, A., Levy, R., Volle, E., & ECOCAPTURE study group. (2025). Behavioral variant frontotemporal dementia as a model for understanding the cognitive and cerebral determinants of verbal creativity. Behavioral and Brain Functions, 21(1), 26. 10.1186/s12993-025-00292-z

Barbot, B. (2018). The Dynamics of Creative Ideation : Introducing a New Assessment Paradigm. Frontiers in Psychology, 9. https://www.frontiersin.org/articles/10.3389/fpsyg.2018.02529

Bartra, O., McGuire, J. T., & Kable, J. W. (2013). The valuation system : A coordinate-based meta-analysis of BOLD fMRI experiments examining neural correlates of subjective value. NeuroImage, 76, 412–427. 10.1016/j.neuroimage.2013.02.063

Basten, U., Biele, G., Heekeren, H. R., & Fiebach, C. J. (2010). How the brain integrates costs and benefits during decision making. Proceedings of the National Academy of Sciences, 107(50), 21767–21772. 10.1073/pnas.0908104107

Battistello, G., Moreno-Rodriguez, S., Volle, E., & Lopez-Persem, A. (2025). Subjective valuation as a domain-general process in creative thinking. Communications Psychology, 3(1), 108. 10.1038/s44271-025-00285-8

Beaty, R. E., Benedek, M., Barry Kaufman, S., & Silvia, P. J. (2015). Default and Executive Network Coupling Supports Creative Idea Production. Scientific Reports, 5(1), Article 1. 10.1038/srep10964

Beaty, R. E., Benedek, M., Silvia, P. J., & Schacter, D. L. (2016). Creative Cognition and Brain Network Dynamics. Trends in Cognitive Sciences, 20(2), 87–95. 10.1016/j.tics.2015.10.004

Beaty, R. E., Benedek, M., Wilkins, R. W., Jauk, E., Fink, A., Silvia, P. J., Hodges, D. A., Koschutnig, K., & Neubauer, A. C. (2014). Creativity and the default network : A functional connectivity analysis of the creative brain at rest. Neuropsychologia, 64, 92–98. 10.1016/j.neuropsychologia.2014.09.019

Beaty, R. E., & Kenett, Y. N. (2023). Associative thinking at the core of creativity. Trends in Cognitive Sciences, 27(7), 671–683. 10.1016/j.tics.2023.04.004

Bendetowicz, D., Urbanski, M., Aichelburg, C., Levy, R., & Volle, E. (2017). Brain morphometry predicts individual creative potential and the ability to combine remote ideas. Cortex; a Journal Devoted to the Study of the Nervous System and Behavior, 86, 216–229. 10.1016/j.cortex.2016.10.021

Bendetowicz, D., Urbanski, M., Garcin, B., Foulon, C., Levy, R., Bréchemier, M.-L., Rosso, C., Thiebaut De Schotten, M., & Volle, E. (2018). Two critical brain networks for generation and combination of remote associations. Brain, 141(1), 217–233. 10.1093/brain/awx294

Benedek, M., Beaty, R. E., Schacter, D. L., & Kenett, Y. N. (2023). The role of memory in creative ideation. Nature Reviews Psychology, 2(4), 246–257. 10.1038/s44159-023-00158-z

Benedek, M., Könen, T., & Neubauer, A. (2012). Associative Abilities Underlying Creativity. Psychology of Aesthetics, Creativity, and the Arts, 6, 273–281. 10.1037/a0027059

Benedek, M., Mühlmann, C., Jauk, E., & Neubauer, A. C. (2013). Assessment of divergent thinking by means of the subjective top-scoring method : Effects of the number of top-ideas and time-on-task on reliability and validity. Psychology of Aesthetics, Creativity, and the Arts, 7(4), 341–349. 10.1037/a0033644

Birney, D. P., Beckmann, J. F., & Seah, Y.-Z. (2016). More than the eye of the beholder : The interplay of person, task, and situation factors in evaluative judgements of creativity. Learning and Individual Differences, 51, 400–408. 10.1016/j.lindif.2015.07.007

Boorman, E. D., Behrens, T. E. J., Woolrich, M. W., & Rushworth, M. F. S. (2009). How green is the grass on the other side? Frontopolar cortex and the evidence in favor of alternative courses of action. Neuron, 62(5), 733–743. 10.1016/j.neuron.2009.05.014

Boorman, E. D., Rushworth, M. F., & Behrens, T. E. (2013). Ventromedial Prefrontal and Anterior Cingulate Cortex Adopt Choice and Default Reference Frames during Sequential Multi-Alternative Choice. The Journal of Neuroscience, 33(6), 2242–2253. 10.1523/jneurosci.3022-12.2013

Ceh, S. M., Edelmann, C., Hofer, G., & Benedek, M. (2022). Assessing Raters : What Factors Predict Discernment in Novice Creativity Raters? The Journal of Creative Behavior, 56(1), 41–54. 10.1002/jocb.515

Chong, T. T.-J., Apps, M., Giehl, K., Sillence, A., Grima, L. L., & Husain, M. (2017). Neurocomputational mechanisms underlying subjective valuation of effort costs. PLoS Biology, 15(2), e1002598. 10.1371/journal.pbio.1002598

Chrysikou, E. G. (2019). Creativity in and out of (cognitive) control. Current Opinion in Behavioral Sciences, 27, 94–99. 10.1016/j.cobeha.2018.09.014

Clairis, N., & Lopez-Persem, A. (2023). Debates on the dorsomedial prefrontal/dorsal anterior cingulate cortex : Insights for future research. Brain: A Journal of Neurology, 146(12), 4826–4844. 10.1093/brain/awad263

De Martino, B., Fleming, S. M., Garrett, N., & Dolan, R. J. (2013). Confidence in value-based choice. Nature Neuroscience, 16(1), 105–110. 10.1038/nn.3279

de Buisonjé, D. R., Ritter, S. M., de Bruin, S., ter Horst, J. M.-L., & Meeldijk, A. (2017). Facilitating creative idea selection : The combined effects of self-affirmation, promotion focus and positive affect. Creativity Research Journal, 29(2), 174–181. 10.1080/10400419.2017.1303308

Diedrich, J., Jauk, E., Silvia, P. J., Gredlein, J. M., Neubauer, A. C., & Benedek, M. (2018). Assessment of real-life creativity : The Inventory of Creative Activities and Achievements (ICAA). Psychology of Aesthetics, Creativity, and the Arts, 12(3), 304–316. 10.1037/aca0000137

Eling, K., Langerak, F., & Griffin, A. (2015). The Performance Effects of Combining Rationality and Intuition in Making Early New Product Idea Evaluation Decisions. Creativity and Innovation Management, 24(3), 464–477. 10.1111/caim.12128

Ellamil, M., Dobson, C., Beeman, M., & Christoff, K. (2012). Evaluative and generative modes of thought during the creative process. NeuroImage, 59(2), 1783–1794. 10.1016/j.neuroimage.2011.08.008

Glimcher, P. W. (2014). Chapter 20—Value-Based Decision Making. In P. W. Glimcher & E. Fehr (Éds.), Neuroeconomics (Second Edition) (p. 373–391). Academic Press. 10.1016/B978-0-12-416008-8.00020-6

Gloy, K., Herrmann, M., & Fehr, T. (2020). Decision making under uncertainty in a quasi realistic binary decision task – An fMRI study. Brain and Cognition, 140, 105549. 10.1016/j.bandc.2020.105549

Grueschow, M., Polania, R., Hare, T. A., & Ruff, C. C. (2015). Automatic versus Choice-Dependent Value Representations in the Human Brain. Neuron, 85(4), 874–885. 10.1016/j.neuron.2014.12.054

Guilford, J. P. (1967). Creativity : Yesterday, Today and Tomorrow. The Journal of Creative Behavior, 1(1), 3–14. 10.1002/j.2162-6057.1967.tb00002.x

Hare, T. A., Malmaud, J., & Rangel, A. (2011). Focusing Attention on the Health Aspects of Foods Changes Value Signals in vmPFC and Improves Dietary Choice. Journal of Neuroscience, 31(30), 11077–11087. 10.1523/JNEUROSCI.6383-10.2011

Hayden, B. Y., Heilbronner, S. R., Pearson, J. M., & Platt, M. L. (2011). Surprise signals in anterior cingulate cortex : Neuronal encoding of unsigned reward prediction errors driving adjustment in behavior. The Journal of Neuroscience: The Official Journal of the Society for Neuroscience, 31(11), 4178–4187. 10.1523/JNEUROSCI.4652-10.2011

Hunt, L. T., Dolan, R. J., & Behrens, T. E. J. (2014). Hierarchical competitions subserving multi-attribute choice. Nature Neuroscience, 17(11), 1613–1622. 10.1038/nn.3836

Hunt, L. T., Kolling, N., Soltani, A., Woolrich, M. W., Rushworth, M. F. S., & Behrens, T. E. J. (2012). Mechanisms underlying cortical activity during value-guided choice. Nature Neuroscience, 15(3), 470–476. 10.1038/nn.3017

Kable, J. W., & Glimcher, P. W. (2009). The Neurobiology of Decision : Consensus and Controversy. Neuron, 63(6), 733–745. 10.1016/j.neuron.2009.09.003

Kahneman, D., & Tversky, A. (1979). Prospect Theory : An Analysis of Decision under Risk. Econometrica, 47(2), 263–291. 10.2307/1914185

Kleinmintz, O. M., Ivancovsky, T., & Shamay-Tsoory, S. G. (2019). The two-fold model of creativity : The neural underpinnings of the generation and evaluation of creative ideas. Current Opinion in Behavioral Sciences, 27, 131–138. 10.1016/j.cobeha.2018.11.004

Kolling, N., Behrens, T. E. J., Mars, R. B., & Rushworth, M. F. S. (2012). Neural mechanisms of foraging. Science (New York, N.Y.), 336(6077), 95–98. 10.1126/science.1216930

Kolling, N., Wittmann, M. K., Behrens, T. E. J., Boorman, E. D., Mars, R. B., & Rushworth, M. F. S. (2016). Value, search, persistence and model updating in anterior cingulate cortex. Nature Neuroscience, 19(10), 1280–1285. 10.1038/nn.4382

Konovalov, A., & Krajbich, I. (2019). Revealed strength of preference : Inference from response times. Judgment and Decision Making, 14(4), 381–394. 10.1017/S1930297500006082

Lazar, M., Miron-Spektor, E., & Mueller, J. S. (2022). Love at first insight : An attachment perspective on early-phase idea selection. Organizational Behavior and Human Decision Processes, 172, 104168. 10.1016/j.obhdp.2022.104168

Levy, D. J., & Glimcher, P. W. (2012). The root of all value : A neural common currency for choice. Current Opinion in Neurobiology, 22(6), 1027–1038. 10.1016/j.conb.2012.06.001

Lin, H., & Vartanian, O. (2018). A Neuroeconomic Framework for Creative Cognition. Perspectives on Psychological Science: A Journal of the Association for Psychological Science, 13(6), 655–677. 10.1177/1745691618794945

Lloyd-Cox, J., Chen, Q., & Beaty, R. E. (2022). The time course of creativity : Multivariate classification of default and executive network contributions to creative cognition over time. Cortex, 156, 90–105. 10.1016/j.cortex.2022.08.008

Lloyd-Cox, J., Pickering, A., & Bhattacharya, J. (2022). Evaluating Creativity : How Idea Context and Rater Personality Affect Considerations of Novelty and Usefulness. Creativity Research Journal, 34(4), 373–390. 10.1080/10400419.2022.2125721

Long, H., & Pang, W. (2015). Rater effects in creativity assessment : A mixed methods investigation. Thinking Skills and Creativity, 15, 13–25. 10.1016/j.tsc.2014.10.004

Lopez-Persem, A., Domenech, P., & Pessiglione, M. (2016). How prior preferences determine decision-making frames and biases in the human brain. eLife, 5, e20317. 10.7554/eLife.20317

Lopez-Persem, A., Moreno-Rodriguez, S., Ovando-Tellez, M., Bieth, T., Guiet, S., Brochard, J., & Volle, E. (2024). How subjective idea valuation energizes and guides creative idea generation. The American Psychologist, 79(3), 403–422. 10.1037/amp0001165

Lopez-Persem, A., Rigoux, L., Bourgeois-Gironde, S., Daunizeau, J., & Pessiglione, M. (2017). Choose, rate or squeeze : Comparison of economic value functions elicited by different behavioral tasks. PLOS Computational Biology, 13(11), e1005848. 10.1371/journal.pcbi.1005848

Mednick, S. (1962). The associative basis of the creative process. Psychological Review, 69(3), 220–232. 10.1037/h0048850

Moreno-Rodriguez, S., Béranger, B., Volle, E., & Lopez-Persem, A. (2025). The human reward system encodes the subjective value of ideas during creative thinking. Communications Biology, 8(1), 1–19. 10.1038/s42003-024-07427-4

Niendam, T. A., Laird, A. R., Ray, K. L., Dean, Y. M., Glahn, D. C., & Carter, C. S. (2012). Meta-analytic evidence for a superordinate cognitive control network subserving diverse executive functions. Cognitive, Affective & Behavioral Neuroscience, 12(2), 241–268. 10.3758/s13415-011-0083-5

Plassmann, H., O’Doherty, J., & Rangel, A. (2007). Orbitofrontal cortex encodes willingness to pay in everyday economic transactions. The Journal of Neuroscience: The Official Journal of the Society for Neuroscience, 27(37), 9984–9988. 10.1523/JNEUROSCI.2131-07.2007

Prabhakaran, R., Green, A. E., & Gray, J. R. (2014). Thin slices of creativity : Using single-word utterances to assess creative cognition. Behavior Research Methods, 46(3), 641–659. 10.3758/s13428-013-0401-7

Quilodran, R., Rothé, M., & Procyk, E. (2008). Behavioral shifts and action valuation in the anterior cingulate cortex. Neuron, 57(2), 314–325. 10.1016/j.neuron.2007.11.031

Rangel, A., & Hare, T. (2010). Neural computations associated with goal-directed choice. Current Opinion in Neurobiology, 20(2), 262–270. 10.1016/j.conb.2010.03.001

Rietzschel, E. F., Nijstad, B. A., & Stroebe, W. (2006). Productivity is not enough : A comparison of interactive and nominal brainstorming groups on idea generation and selection. Journal of Experimental Social Psychology, 42(2), 244–251. 10.1016/j.jesp.2005.04.005

Rietzschel, E. F., Nijstad, B. A., & Stroebe, W. (2010). The selection of creative ideas after individual idea generation : Choosing between creativity and impact. British Journal of Psychology, 101(1), 47–68. 10.1348/000712609X414204

Runco, M. A., & Jaeger, G. J. (2012). The Standard Definition of Creativity. Creativity Research Journal, 24(1), 92–96. 10.1080/10400419.2012.650092

Rushworth, M. F. S., Kolling, N., Sallet, J., & Mars, R. B. (2012). Valuation and decision-making in frontal cortex : One or many serial or parallel systems? Current Opinion in Neurobiology, 22(6), 946–955. 10.1016/j.conb.2012.04.011

Samuelson, P. A. (1937). A Note on Measurement of Utility. The Review of Economic Studies, 4(2), 155–161. 10.2307/2967612

Shenhav, A., & Botvinick, M. (2015). Uncovering a missing link in anterior cingulate research. Neuron, 85(3), 455–457. 10.1016/j.neuron.2015.01.020

Shenhav, A., Botvinick, M. M., & Cohen, J. D. (2013). The expected value of control : An integrative theory of anterior cingulate cortex function. Neuron, 79(2), 217–240. 10.1016/j.neuron.2013.07.007

Shi, L., Sun, J., Xia, Y., Ren, Z., Chen, Q., Wei, D., Yang, W., & Qiu, J. (2018). Large-scale brain network connectivity underlying creativity in resting-state and task fMRI : Cooperation between default network and frontal-parietal network. Biological Psychology, 135, 102–111. 10.1016/j.biopsycho.2018.03.005

Sowden, P., Pringle, A., & Gabora, L. (2015). The shifting sands of creative thinking : Connections to dual-process theory. Thinking & Reasoning, 21, 40–60. 10.1080/13546783.2014.885464

Sullivan, N. J., & Huettel, S. A. (2021). Healthful choices depend on the latency and rate of information accumulation. Nature Human Behaviour, 5(12), 1698–1706. 10.1038/s41562-021-01154-0

Tsetsos, K., Wyart, V., Shorkey, S. P., & Summerfield, C. (2014). Neural mechanisms of economic commitment in the human medial prefrontal cortex. eLife, 3, e03701. 10.7554/eLife.03701

Volle, E. (2018). Associative and controlled cognition in divergent thinking : Theoretical, experimental, neuroimaging evidence, and new directions. In The Cambridge handbook of the neuroscience of creativity (p. 333–360). Cambridge University Press. 10.1017/9781316556238.020

Volz, K. G., Schubotz, R. I., & von Cramon, D. Y. (2005). Frontomedian activation depends on both feedback validity and valence : fMRI evidence for contextual feedback evaluation. NeuroImage, 27(3), 564–571. 10.1016/j.neuroimage.2005.04.026

Wang, X., Chen, Q., Zhuang, K., Zhang, J., Cortes, R. A., Holzman, D. D., Fan, L., Liu, C., Sun, J., Li, X., Li, Y., Feng, Q., Chen, H., Feng, T., Lei, X., He, Q., Green, A. E., & Qiu, J. (2024). Semantic associative abilities and executive control functions predict novelty and appropriateness of idea generation. Communications Biology, 7(1), 703. 10.1038/s42003-024-06405-0

Wunderlich, K., Rangel, A., & O’Doherty, J. P. (2009). Neural computations underlying action-based decision making in the human brain. Proceedings of the National Academy of Sciences, 106(40), 17199–17204. 10.1073/pnas.0901077106

Wunderlich, K., Rangel, A., & O’Doherty, J. P. (2010). Economic choices can be made using only stimulus values. Proceedings of the National Academy of Sciences of the United States of America, 107(34), 15005–15010. 10.1073/pnas.1002258107

Yeo, B. T. T., Krienen, F. M., Sepulcre, J., Sabuncu, M. R., Lashkari, D., Hollinshead, M., Roffman, J. L., Smoller, J. W., Zöllei, L., Polimeni, J. R., Fischl, B., Liu, H., & Buckner, R. L. (2011). The organization of the human cerebral cortex estimated by intrinsic functional connectivity. Journal of Neurophysiology, 106(3), 1125–1165. 10.1152/jn.00338.2011

Zabelina, D. L., & Andrews-Hanna, J. R. (2016). Dynamic network interactions supporting internally-oriented cognition. Current Opinion in Neurobiology, Systems neuroscience, 40, 86–93. 10.1016/j.conb.2016.06.014

Zhu, Y., Ritter, S. M., & Dijksterhuis, A. (2021). The effect of rank-ordering strategy on creative idea selection performance. European Journal of Social Psychology, 51(2), 360–376. 10.1002/ejsp.2743

Zhu, Y., Ritter, S. M., Müller, B. C. N., & Dijksterhuis, A. (2017). Creativity : Intuitive processing outperforms deliberative processing in creative idea selection. Journal of Experimental Social Psychology, 73, 180–188. 10.1016/j.jesp.2017.06.009

