## Supplemental for "Neural investigation of idea selection during creative thinking reveals value-based mechanisms"

### Supplementary Results

*Behavioral results: model comparison*

All choice models were based on a classical function from the decision-making literature: the softmax function (see Equation A in Methods). The softmax function explains the probability of choosing a specific idea in function of the ideas' DV. Here, it explains the probability of choosing the FGAT-distant response in function of the DV between the FGAT-distant and FGAT-first response (DV = $V_{FGAT\text{-}distant}$-$V_{FGAT\text{-}first}$). Importantly, the compared models (see their equations in Methods) differed in how the DVs were computed ("value-before-choice" versus "value-during-choice" models) and whether a constant was added to the DVs.

First, models varied in their computation of DVs, specifically in the parameters used to compute values. Indeed, previous studies from the lab demonstrated that, during idea evaluation, individuals assign subjective values to ideas as a function of their originality and adequacy ratings; specifically, a 'Constant Elasticity of Substitution' function (CES, see Equation B in Methods). By fitting this function to individuals' ratings, we estimate two valuation parameters: $\alpha_{rating}$ and $\delta_{rating}$. $\alpha_{rating}$ captures the weight given to originality relative to adequacy in likeability ratings (how much originality ratings explains the variability of likeability ratings relative to adequacy). If $\alpha_{rating}$ is greater than 0.5, likeability is more driven by originality (and vice versa). $\delta_{rating}$ captures the preference for an equilibrium between originality and adequacy. If $\delta_{rating}$ is lower than 1, likeability increases when both adequacy and originality are balanced. If $\delta_{rating}$ is greater than 1, likeability increases when one dimension outweighs the other. In our model comparison, the "value-before-choice" models, i.e. models 1 and 2, featured values computed using the valuation parameters previously estimated from the CES function using likeability ratings ($\alpha_{rating}$ and $\delta_{rating}$) as fixed parameters. "Value-before-choice" models, i.e., models 3 and 4, featured values computed using free parameters ($\alpha_{choice}$ and $\delta_{choice}$, estimated during the softmax model fitting). Second, besides DV computation, the models also differed in whether they included a constant $\gamma$, a free parameter estimated during fitting which captures an additive bias against ($\gamma < 0$) or towards ($\gamma > 0$) FGAT-distant ideas (Lopez-Persem et al., 2016). Only models 2 and 4 included $\gamma$. Note that, in all models, the softmax function also includes a free parameter $\beta$ (the inverse temperature that captures the expected stochasticity, or incoherence, in individuals' choices).

The model comparison revealed that the model that best explained choices was a "value-during-choice" model, i.e. model 4, which included the $\gamma$ parameter (additive bias), and used the free $\alpha_{choice}$ and $\delta_{choice}$ parameters, rather than the fixed $\alpha_{rating}$ and $\delta_{rating}$ parameters (model 1: estimated model frequency (Ef) = 0.002, exceedance probabilities (Xp) = 0; model 2: Ef = 0.034, Xp = 0; model 3: Ef = 0.005, Xp = 0; model 4: Ef = 0.96, Xp = 1; model 4's balanced accuracy range: [0.49 ; 0.89], Figure 1A). This suggests that the values that drive choices slightly differ from those expressed during likeability ratings. Next, we investigated model 4's parameters.

*Behavioral results: valuation parameter estimation and comparison*

First, we found that, on average, participants put less weight on originality than on adequacy ($\alpha_{choice}$ = 0.38±0.02 (M±SEM), one-sample two-tailed t-test against 0.5: $t(107) = -5.57$, $p < 0.001$). We compared the $\alpha_{choice}$ parameter to the $\alpha_{rating}$ parameter and found that during choices, individuals' weighting of originality had dropped, compared to during the likeability rating task ($\alpha_{rating}$ = 0.44±0.02 (M±SEM), $\alpha_{rating}$-$\alpha_{choice}$ = 0.07±0.01 (M±SEM), one-sample two-tailed t-test: $t(107) = 4.12$, $p < 0.001$).

Then, we found that the $\delta_{choice}$ parameter was not significantly different from 1, suggesting that there was no group preference for ideas with an equilibrium in originality and adequacy over ideas that are more extreme on one or the other dimension ($\delta_{choice}$ = 0.84±0.09 (M±SEM), one-sample two-tailed t-test against 1: t(107) = -1.71, p = 0.09). We compared the $\delta_{choice}$ parameter to the $\delta_{rating}$ parameter and found that, during choices, participants' preference for equilibrium had waned, compared to during the likeability rating task ($\delta_{rating}$ = 0.49±0.14 (M±SEM); $\delta_{rating}$-$\delta_{choice}$ = -0.34±0.14 (M±SEM), one-sample two-tailed t-test: t(107) = -2.47, p = 0.02).

Importantly, the valuation parameters estimated from the choice task and the rating task were correlated (correlation between $\alpha_{rating}$ and $\alpha_{choice}$: r = 0.74, p < 0.001; $\delta_{rating}$ and $\delta_{choice}$: r = 0.35, p < 0.001; Figure 1B), meaning that, despite slight differences, preferences stayed coherent within individuals.

Next, we found that the $\gamma$ parameter was significantly higher than 0, suggesting that at the group level, there was an additive bias for FGAT-distant ideas, against FGAT-first ideas ($\gamma$ = 0.38±0.18 (M±SEM), one-sample two-tailed t-test: t(107) = -2.14, p = 0.03). In other words, FGAT-distant responses typically received a small value bonus when compared to FGAT-first responses. This aligns with the choice task instructions (i.e., to choose the response they would have preferred to provide in the FGAT-distant condition). Interestingly, it seemed paradoxical that the $\gamma$ parameter would be in favor of FGAT-distant ideas (which are more original ideas than in FGAT-first) while $\alpha_{choice}$ was in favor of adequacy. We wondered whether this reflected a coping mechanism, where participants who overvalued adequacy attributed an added value to FGAT-distant ideas to cancel this out. On the contrary, we found that the participants who put a higher weight on adequacy during choices (lower $\alpha_{choice}$) had a lower additive bias in favor of FGAT-distant ideas (lower $\gamma$) (correlation between $\alpha_{choice}$ and $\gamma$ parameter : r = 0.59; p < 0.001).

*Behavioral results: correlation between valuation parameters and creative performance*

We sought to confirm that the decision values computed during choices (using $\alpha_{choice}$ and $\delta_{choice}$ as valuation parameters) were relevant to creative performance: since our past work showed that individuals' $\alpha_{rating}$ and $\delta_{rating}$ predicted their creativity scores in the battery of creativity tests, we tested whether $\alpha_{choice}$ and $\delta_{choice}$ (as well as $\gamma$) showed similar patterns. First, we found that the number of unique answers in FGAT-distant correlated with $\alpha_{choice}$ (r = 0.30; p = 0.002, Figure 1E) and with $\delta_{choice}$ (r = -0.25; p = 0.007, Figure 1F), but not with the $\gamma$ parameter (see Table S2).

Then, to test the generalizability of this result, we used the battery of external creativity tests to measure creative performance in other domains of creativity (in the Drawing Task, the Inventory of Creative Activities and Achievements (ICAA), the Alternative Uses Task (AUT), the Combination of Associates Task (CAT) and the Associative Fluency Task, see Methods for details). We performed a principal component analysis (PCA) to reduce the dimensions of external creativity measures and found two principal components with an eigenvalue >1 (PC1 and PC2, see Methods for more details). PC1 seemed to capture associative abilities and real-life creative abilities, while PC2 gathered convergent and divergent thinking abilities. In line with the previous results using FGAT-distant creative performance, we found that participants favoring originality during creative idea comparison (higher $\alpha_{choice}$) yielded higher creativity scores (correlation between $\alpha_{choice}$ and PC2: r = 0.26; p = 0.007, Figure 1G). Note that $\alpha_{choice}$ did not significantly correlate with PC1, and that $\delta_{choice}$ and the $\gamma$ parameter did not significantly correlate with PC1 or PC2 (see Table S2).

| Kaiser-Meyer-Olkin (KMO) measure of sampling adequacy | | | |
|---|---|---|---|
| task | sampling adequacy | | |
| general | 0.575 | | |
| drawing task | 0.574 | | |
| ICAA | 0.587 | | |
| AUT | 0.550 | | |
| CAT | 0.451 | | |
| associative fluency task | 0.640 | | |
| **Explained Variance** | | | |
| component | eigenvalue | variance % | cumulated variance % |
| PC1 | 1.87 | 37.5% | 37.5% |
| PC2 | 1.15 | 23% | 60.5% |
| PC3 | 0.9 | 18.2% | 78.7% |
| PC4 | 0.58 | 11.5% | 90.2% |
| PC5 | 0.49 | 9.76% | 100% |
| **Component contributions** | | | |
| task | | loadings on PC1 | loadings on PC2 |
| drawing task | | 0.47 | 0.61 |
| ICAA | | 0.84 | -0.08 |
| AUT | | -0.08 | 0.57 |
| CAT | | -0.15 | 0.80 |
| associative fluency task | | 0.80 | 0.01 |

**Table S1 : Principal Components Analysis**

| | number of unique FGAT-distant responses | PC1 | PC2 |
|---|---|---|---|
| $\alpha$ choice | r = 0.31 ; p = 0.0011 | r = -0.02 ; p = 0.81 | r = 0.23 ; p = 0.02 |
| $\delta$ choice | r = -0.26 ; p = 0.007 | r = 0.04 ; p = 0.65 | r = -0.07 ; p = 0.45 |
| $\gamma$ | r = 0.15 ; p = 0.13 | r = -0.05 ; p = 0.59 | r = 0.11 ; p = 0.28 |

**Table S2 : All correlations between creative performance and the parameters estimated using the choice task ($\alpha_{choice}$, $\delta_{choice}$ and the $\gamma$ parameter)**

*Behavioral results: Control analyses of the relationship between $RT_{FGAT\text{-}distant}$ and DV*

While the current study reports a negative relationship between *$RT_{FGAT\text{-}distant}$ and DV,* comparable results were found in previous studies from the lab (Lopez-Persem et al., 2024; Moreno-Rodriguez et al., 2025) where $RT_{FGAT\text{-}distant}$ decreased with the value of the FGAT-distant option. Given that DVs in the current analysis contain the value of the FGAT-distant option, we performed control analyses to make sure the current results were not just explained by the value of the FGAT-distant, but indeed by the difference of the value of the FGAT-distant and of the FGAT-first option.

Using the Matlab VBA toolbox (c.f. the Model comparison section in the Methods), we performed a model comparison analysis, comparing two models: a linear model of $RT_{FGAT\text{-}distant}$ explained by DV (model 1), and a linear model of $RT_{FGAT\text{-}distant}$ explained by $V_{FGAT\text{-}distant}$ (model 2). Confirming the current results reported in the main text, this model comparison revealed that the variable that best explained

$RT_{FGAT\text{-}distant}$ was DV (model 1: estimated model frequency (Ef) = 0.37, exceedance probabilities (Xp) = 0; fit $r^2$ = 0.037; model 2: Ef = 0.63, Xp = 1; fit $r^2$ = 0.048).

*Neuroimaging results: DV encoding during the FGAT-distant using FIR analyses locked on cue-onset*

When investigating the encoding of DV in the ROIs during the FGAT, we performed FIR analyses locked on response onset (reported in the main text) and locked on cue onset, reported here: we found that the correlation between DVs and the activity within the vmPFC ROI was not significantly different from 0, at any time point of the analysis time window locked on cue onset (Figure S1A, see Table S3 for all results). The analysis time window was *a priori* defined as the time window approximating the mean cue-to-response interval in FGAT-distant trials ($RT_{FGAT\text{-}distant}$ = 6.7 ± 0.4 seconds, M±SEM). We investigated whether the correlation was significant over the entire analysis time window, and found a non-significant correlation between DVs and the activity within the vmPFC ROI (see Table S3 for all results). Looking at the encoding of the value of the FGAT-first and FGAT-distant responses individually, we found a negative encoding of the value of the FGAT-first response at 3 time points near the response onset (Figure S1B, see Table S3 for all results). Again, we investigated whether it was significant over the entire analysis time window and found a significant negative correlation. There was no significant correlation between the value of the FGAT-distant response and the activity within the vmPFC ROI (see Table S3 for all results).

In parallel, we found that the correlation between DVs and the activity within the dACC ROI was not significantly different from 0 at any time point (Figure S1C, see Table S3 for all results). We investigated whether the correlation was significant in the analysis time window, and found a non-significant correlation between DVs and the activity within the dACC ROI (see Table S3 for all results). Looking at the encoding of the value of the FGAT-first and FGAT-distant responses individually, we found non-significant results at all time points, and in the analysis time window (see Table S3 for all results).

Together, these non-significant results in analyses locked on cue onset confirm the necessity to use FIR analyses and lock analyses around the response onset, which tackles the dilution of results due to different response times found when using classical HRF analyses.

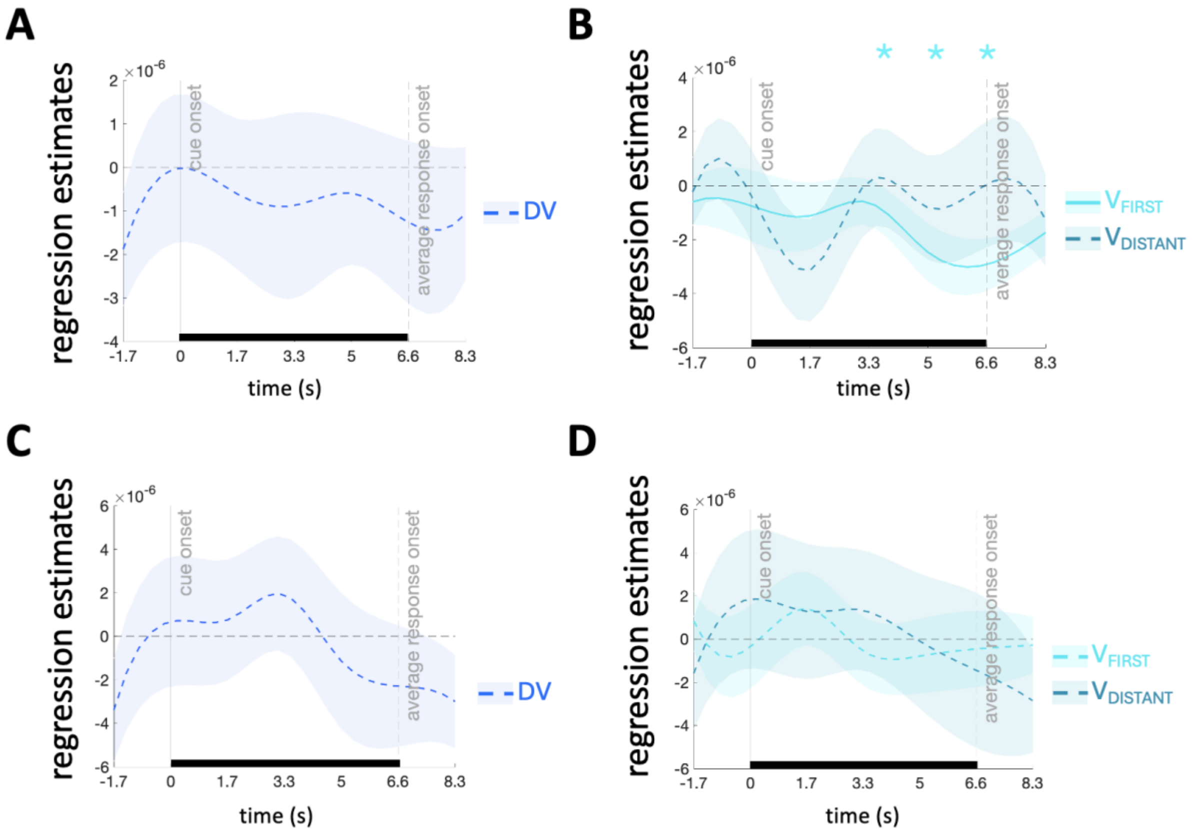


**Figure S1: Time course regression estimates for the decision value and for the values of FGAT-fist and FGAT-distant responses in the vmPFC and dACC ROIs locked on cue onset**

**(A)** Time course of the peri-cue regression estimate for decision value (DV) during the FGAT-distant task, in the vmPFC ROI (shown in Figure 4A) **(B)** Time course of the peri-cue regression estimate for the values of the FGAT-first and FGAT-distant responses during the FGAT-distant task, in the vmPFC ROI (shown in Figure 4A) **(C)** Time course of the peri-cue regression estimate for decision value (DV) during the FGAT-distant task, in the dACC ROI (shown in Figure 4B) **(D)** Time course of the peri-cue regression estimate for the values of the FGAT-first and FGAT-distant responses during the FGAT-distant task, in the dACC ROI (shown in Figure 4B). The shaded areas represent the SEM. Solid lines indicate a significant correlation ($p<0.05$) during the analysis time window (represented here with a horizontal black line), dashed lines indicate non-significant ($p>0.05$) correlation during the analysis time window. The analysis time window is an *a priori* defined time window approximating the mean cue-to-response interval in FGAT-distant trials ($RT_{FGAT\text{-}distant} = 6.7 \pm 0.4$ seconds, M±SEM). Results in bold ($p<0.05$) are significant. * indicate significant ($p<0.05$) correlations at specific time points. n=38.

| variable | time-lock | ROI | time relative to response onset (s) | β (M±SEM) (x10-5) | t-stat(37) | p-value |
|---|---|---|---|---|---|---|
| positive decision value | cue onset | vmPFC | analysis time window | -0.07±0.02 | -0.64 | 0.74 |
| | | | 0 | -0.06±0.03 | -0.35 | 0.637 |
| | | | 1.66 | -0.09±0.03 | -0.41 | 0.658 |
| | | | 3.32 | -0.06±0.03 | -0.36 | 0.639 |
| | | | 4.98 | -0.13±0.03 | -0.67 | 0.748 |
| | | | 6.64 | -0.11±0.02 | -0.70 | 0.755 |
| FGAT-first value | cue onset | vmPFC | **analysis time window** | **-0.16±0.01** | **-2.68** | **0.006** |
| | | | 0 | -0.11±0.02 | -0.90 | 0.186 |
| | | | 1.66 | -0.06±0.01 | -0.73 | 0.235 |
| | | | **3.32** | **-0.25±0.02** | **-2.29** | **0.014** |
| | | | **4.98** | **-0.29±0.02** | **-2.99** | **0.003** |
| | | | **6.64** | **-0.17±0.01** | **-2.11** | **0.021** |
| FGAT-distant value | cue onset | vmPFC | analysis time window | -0.09±0.02 | -0.70 | 0.756 |
| | | | 0 | -0.31±0.03 | -1.65 | 0.946 |
| | | | 1.66 | 0.02±0.03 | 0.10 | 0.459 |
| | | | 3.32 | -0.08±0.03 | -0.42 | 0.663 |
| | | | 4.98 | 0.00±0.04 | 0.02 | 0.492 |
| | | | 6.64 | -0.13±0.03 | -0.75 | 0.772 |
| negative decision value | cue onset | dACC | analysis time window | 0.05±0.03 | 0.26 | 0.399 |
| | | | 0 | -0.08±0.05 | -0.27 | 0.606 |
| | | | 1.66 | -0.19±0.04 | -0.73 | 0.764 |
| | | | 3.32 | 0.11±0.05 | 0.36 | 0.359 |
| | | | 4.98 | 0.23±0.05 | 0.82 | 0.209 |
| | | | 6.64 | 0.30±0.03 | 1.40 | 0.085 |
| FGAT-first value | cue onset | dACC | analysis time window | -0.02±0.02 | -0.14 | 0.554 |
| | | | 0 | 0.14±0.03 | 0.83 | 0.205 |
| | | | 1.66 | -0.06±0.03 | -0.35 | 0.637 |
| | | | 3.32 | -0.08±0.03 | -0.46 | 0.674 |
| | | | 4.98 | -0.05±0.03 | -0.26 | 0.602 |
| | | | 6.64 | -0.03±0.02 | -0.22 | 0.586 |
| FGAT-distant value | cue onset | dACC | analysis time window | 0.00±0.00 | 0.00 | 0.501 |
| | | | 0 | 0.13±0.05 | 0.41 | 0.657 |
| | | | 1.66 | 0.13±0.05 | 0.47 | 0.681 |
| | | | 3.32 | -0.01±0.05 | -0.03 | 0.488 |
| | | | 4.98 | -0.14±0.05 | -0.40 | 0.345 |
| | | | 6.64 | -0.28±0.04 | -1.18 | 0.123 |

**Table S3: Time-point-wise FIR statistics for parametric modulation in the vmPFC and dACC ROIs during the FGAT-distant, locked on cue onset.**

The “analysis time window” is an *a priori* defined time window approximating the mean cue-to-response interval in FGAT-distant trials ($RT_{FGAT\text{-}distant}$ = 6.7 ± 0.4 seconds, M±SEM). Results in bold are significant ($p<0.05$)

**Supplementary Methods**

*FGAT task instructions*

In the FGAT, we divided instructions into successive slides, illustrated with screenshots of what the task would look like, and participants read them at their own pace. The instructions were as follows (translated from French):
*FGAT-first:* "In each trial, you will see a word appear on the screen. You will have to state the first word that comes to your mind without thinking and as fast as possible. As soon as you have a word in mind, you will need to press your index finger to give your response to the experimenter. The word "Response?" will then appear on the screen: you can say the word out loud to the experimenter. The experimenter will write down the word heard. You must verify that it is indeed the word you said. If yes, press with your ring finger to confirm your answer. If not, press with your index finger to repeat your response. Please note that this is only to confirm that the word has been heard correctly. You do not have the right to change your mind. Once your answer is confirmed, a new word will appear on the screen; proceed in the same way. Be aware! You will have 10 seconds to think of your response. If you take too long to find an answer, we will move on to the next trial. Before pressing with your index finger, wait until you have an answer in mind: you must say it directly to the experimenter without taking additional time to think. Conjugated verbs, proper nouns, and groups of words are not accepted."
*FGAT-distant:* "Similar to the previous task, with each new trial, you will see a word appear on the screen. This time, you must respond with a word that is not typically associated with the displayed word but is still somehow related. Think creatively: the association between the displayed word and your response should be original, unusual, surprising, while still being understandable to someone else. As soon as you have a word in mind, you will need to press your index finger to give your response to the experimenter. The word "Response?" will then appear on the screen: you can say the word out loud to the experimenter. The experimenter will write down the word heard. You must verify that it is indeed the word you said. If yes, press with your ring finger to confirm your answer. If not, press with your index finger to repeat your response. Please note that this is only to confirm that the word has been heard correctly. You do not have the right to change your mind. Once your answer is confirmed, a new word will appear on the screen; proceed in the same way. Be aware! You will have 20 seconds to think of your response. If you take too long to find an answer, we will move on to the next trial. Before pressing with your index finger, wait until you have an answer in mind: you must say it directly to the experimenter without taking additional time to think. Conjugated verbs, proper nouns, and groups of words are not accepted."

*Rating task instructions*

For the likeability rating task, we divided instructions into successive slides, illustrated with screenshots of what the task would look like, and participants read them at their own pace. The instructions were as follows (translated from French):
"As a reminder, at the beginning of this experiment, you performed two tasks. In response to a word (for example, "mother"), you were required to give two types of responses: the first response that comes to your mind (for example, "mother-father") and an unusual but understandable response (for example, "mother-nature"). In this task, with each new trial, you will see a pair of words on the screen. The first word is part of the words you have seen in the previous tasks. The second word is a word possibly associated with the first word, which can be a word you proposed in the previous tasks or another suggestion. The task is to indicate how much you like the association between these two words in the context of the previous task (where you had to provide an unusual response). For example, how much

would you have liked to respond with the word "father" to the word "mother" in the previous task? Below the association, an evaluation scale will appear, with a heart. Using this scale, you will indicate how much you like this association. Move the cursor using your index and middle fingers. Confirm your response by pressing with your ring finger. Be aware! You must move the cursor, even if you are perfectly neutral (in this case, move and return the cursor to the middle). Use the entire scale: do not always respond the same way! Reminder: Indicate how much you would have liked to propose the second word of the association in response to the previous task (unusual response). If it's a response you have given before, indicate how satisfied you are with that response."

For the originality and adequacy rating task, we divided instructions into successive slides, illustrated with screenshots of what the task would look like, and participants read them at their own pace. The instructions were as follows (translated from French):
"As a reminder, at the beginning of this experiment, you performed two tasks. In response to a word (for example, "mother"), you were required to give two types of responses: the first response that comes to your mind (for example, "mother-father") and an unusual but understandable response (for example, "mother-nature"). Similar to the previous task, with each new trial, you will see a pair of words on the screen. You will evaluate this word association based on two specific criteria. Firstly, does this association seem appropriate to you, meaning understandable, relevant, fitting? To respond, a relevance evaluation scale will appear. Move the cursor using the "left arrow" and "right arrow" keys. Confirm your response by pressing "space." If the association seems entirely relevant to you, move the cursor to the right. If you do not see a connection between the words, move the cursor to the left. Any evaluation between these two extremes is possible. Once your response is confirmed, you will evaluate a second criterion, that of the originality of this word association: how unusual and surprising you find this association. For this, a second evaluation scale will appear. This scale will allow you to indicate how original you think the association is. Move the cursor using the "left arrow" and "right arrow" keys. Confirm your response by pressing "space." Be aware! You must move the cursor, even if you are perfectly neutral (in this case, move and return the cursor to the middle). Try to use the entire scale: do not always respond the same way."

*Rating task trials: selection of the FGAT associations for the rating tasks*

The number of trials in the rating task varied between participants as they saw only associations that followed these criteria:

- the first three letters of the FGAT-first and FGAT-distant responses to the cue word must be different
- one of the responses must not contain the other (e.g. 'count' and 'accountant' have the word 'count' in common and would not respect this criterion)
- the FGAT-first and FGAT-distant responses to one cue word must be comparable in length (no more than five letters of difference)
- the FGAT-distant response must not be too long (no more than nine letters)

Then, we randomly selected one-fifth of the accepted cue words and paired them with five other responses that the participant had not generated: four of those from a different study's dataset, including (i) a highly frequent FGAT-first response, (ii) a highly infrequent FGAT-first response, (iii) a highly frequent FGAT-distant response and (iv) a highly infrequent FGAT-distant response. The fifth response was an unrelated word (for example, the cue word 'cow' was associated with the unrelated response 'inverse'). Overall, this added up to a maximum of 186 rating trials (less if the 124 FGAT-first and

FGAT-distant associations did not all follow the four criteria mentioned above), selected for their diversity and randomized before the start of the task.

*Choice task instructions*

For the choice task, we divided instructions into successive slides, illustrated with screenshots of what the task would look like, and participants read them at their own pace. The instructions were as follows (translated from French):
“As a reminder, at the beginning of this experiment, you performed two tasks. In response to a word (for example, "mother"), you were required to give two types of responses: the first response that comes to your mind (for example, "mother-father") and an unusual but understandable response (for example, "mother-nature"). Similar to the previous task, with each new trial, you will see a word at the top of the screen (for example, "mother"), but this time it will be associated with two other proposals (for example, "father" and "nature"). Choose the association you prefer: "mother" with "father" or "mother" with "nature." In other words, would you have preferred to respond with "father" or "nature" to the word "mother" in the second task (unusual response)? To select a response, use [in the MRI] your index finger (left response) or your middle finger (right response) / [outside the MRI] the left arrow (left response) or the right arrow (right response) of your keyboard. You do not need to validate your choice; it will be framed and validated as soon as you press. Another trial will then start.”
